# A Computational Modelling Approach for Deriving Biomarkers to Predict Cancer Risk in Premalignant Disease

**DOI:** 10.1101/020222

**Authors:** Andrew Dhawan, Trevor A. Graham, Alexander G. Fletcher

## Abstract

The lack of effective biomarkers for predicting cancer risk in premalignant disease is a major clinical problem. There is a near-limitless list of candidate biomarkers and it remains unclear how best to sample the tissue in space and time. Practical constraints mean that only a few of these candidate biomarker strategies can be evaluated empirically and there is no framework to determine which of the plethora of possibilities is the most promising. Here we have sought to solve this problem by developing a theoretical platform for *in silico* biomarker development. We construct a simple computational model of carcinogenesis in premalignant disease and use the model to evaluate an extensive list of tissue sampling strategies and different molecular measures of these samples. Our model predicts that: (i) taking more biopsies improves prognostication, but with diminishing returns for each additional biopsy; (ii) longitudinally-collected biopsies provide slightly more prognostic information than a single biopsy collected at the latest possible time-point; (iii) measurements of clonal diversity are more prognostic than measurements of the presence or absence of a particular abnormality and are particularly robust to confounding by tissue sampling; and (iv) the spatial pattern of clonal expansions is a particularly prognostic measure. This study demonstrates how the use of a mechanistic framework provided by computational modelling can diminish empirical constraints on biomarker development.

## Introduction

Each year, tens of thousands of patients in the UK are diagnosed with a premalignant disease, a benign condition that predisposes to the future develop of cancer. Examples of common premalignant diseases include Barrett’s Oesophagus [1], Ductal Carcinoma *in situ* (DCIS) of the breast [2], benign prostatic intraepithelial neoplasia (PIN) [3], and carcinoma *in situ* in the bladder [4]. The clinical management of patients with premalignant disease is a major challenge: in order to prevent cancer, patients are typically enrolled into longitudinal screening programmes that aim to detect (and then treat) patients who show early signs of progression to cancer. However, while having a premalignant disease increases the *average* risk of developing cancer compared to the unaffected population, the cancer risk for any individual is highly variable and generally quite low. For example, patients with Barrett’s Oesophagus have an average 40-fold increased lifetime risk of developing adenocarcinoma, but the progression rate per patient per year is less than 0.5% [5] and so many of these patients will not progress to cancer in their lifetime. As a result, it is arguable that surveying an average (low-risk) patient is unnecessary as they are unlikely to ever progress to cancer. In addition, the surveillance process is typically unpleasant for the patient, and is very costly to health-care providers. In view of these facts together, premalignant disease is often described as both *over-diagnosed* and *over-treated* [6], and consequently there is a pressing clinical need to be able to accurately stratify cancer risk in these patients.

Prognostic biomarkers are central to current risk-stratification strategies. Here a biomarker is defined as an analysable property of the diseased tissue that correlates with the risk of progressing to cancer. In general, it remains unclear which of the plethora of potential biological features that could be assayed (morphological, gene expression, mutation, or other features) offers the most potential for prognostic value. Pathological grading and staging remain the most widespread biomarkers in current use; these biomarkers are descriptions of the morphological features of the disease. The current state-of-the-art biomarkers are molecular in nature, and typically quantify the aberrant expression of a panel of carefully-chosen genes. For example, the Oncotype DX assay analyses the activity of 21 genes to determine a score quantifying risk of recurrent breast cancer and response to chemotherapy [7]. Genetically based biomarkers include EGFR mutations in non-small cell lung cancer [8] and *TP53* abnormalities in Barrett’s Oesophagus [9]. The limited predictive value of existing biomarkers has prevented their widespread clinical use [10], and for many diseases such as DCIS [11] and inflammatory bowel disease [12] no prognostic biomarkers have yet been identified.

All biomarkers require the diseased tissue to be sampled. Needle biopsies are the predominant sampling method, although other tissue collection methods such as endoscopic brushings or cell washings are sometimes used. However, typically the prognostic optimality of different sampling schemes, including whether samples should be collected longitudinally, has not been evaluated. Furthermore, given the fact that taking a biopsy is an invasive procedure, an empirical evaluation of different tissue sampling schemes is largely unfeasible.

Cancer development is fundamentally an evolutionary process: the acquisition of random somatic mutations can cause a cell to develop an evolutionary advantage over its neighbours, and so drive the clonal expansion of the mutant. Repeated rounds of mutation and clonal selection can lead to the development of a malignant tumour. When viewed from this evolutionary perspective, a biomarker may be thought of as a predictor of the *evolutionary trajectory* of the disease; a successful biomarker is one that sensitively and specifically detects which premalignant lesions are (rapidly) evolving towards cancer. However, existing biomarker development efforts do not explicitly consider the evolutionary process they seek to assay, instead relying on the identification of a small set of genes that are aberrantly expressed in high-risk cases [10]. The recent appreciation that carcinogenesis is a highly stochastic process [13], in which many different combinations of genetic alterations and gene expression changes contribute to the same malignant phenotypes, has led to doubts about the utility of such “candidate gene” approaches [14]. Alternative biomarker development strategies attempt to assay the underlying evolutionary *process* itself. Quantification of within-tumour diversity, as a proxy measure of the probability that the tumour has evolved a well-adapted “dangerous” clone, is one such measure that has shown efficacy in a variety of cancer types [15, 16, 17]. Whilst most studies have focused on the quantification of within-tumour genetic diversity, it is noteworthy that quantification of phenotypic heterogeneity also shows prognostic value [18, 19].

Mathematical models are tools that have the potential to diminish the inherent constraints of empirical biomarker development. Due to the relative ease with which a mathematical model of cancer evolution can be analysed, potentially exhaustive searches of candidate biomarkers can be performed *in silico.* This is the idea that we develop in this study.

Mathematical modelling has a rich history in cancer research, and is increasingly used as a tool to investigate and test hypothesized mechanisms underlying tumour evolution [20]. A common approach is to consider spatially homogeneous well-mixed populations [21], using multi-type Moran models of constant or exponentially growing size [22] or multi-type branching processes [23]. Other work has highlighted the impact of spatial dynamics on the evolutionary process [24]. More complex models have coupled a discrete representation of the movement and proliferation of individual cells to a continuum description of microenvironment factors such as oxygen concentration and extracellular matrix composition. Such models, in particular the pioneering work of Anderson and colleagues [25, 26], demonstrate the significant selective force imposed by microenvironmental conditions such as hypoxia. A recent discussion of the use of ecological and evolutionary approaches to study cancer is provided by Korolev et al. [27]. The majority of models of tumour evolution have focused on the rates of invasion and accumulation of mutations, and how these depend on factors such as modes of cell division and spatial heterogeneity in cell proliferation and death. Defining statistics that correlate with prognosis in these kinds of models is an unaddressed problem.

Here we use mathematical modelling as a novel platform for *in silico* biomarker development. We develop a simple mathematical model of tumour evolution, and use the model to evaluate the prognostic value of a range of different potential biomarker measures and different tissue sampling schemes.

## Materials and methods

### Computational model of within-tumour evolution and biopsy sampling

To simulate the growth and dynamics of a pre-cancerous lesion, we consider a continuous-time spatial Moran process model of clonal evolution [28] on a two-dimensional square lattice, which may be thought of as a mathematical representation of an epithelial tissue. This description is similar to a model of field cancerization proposed by Foo et al. [29], although our model differs in several respects, which we describe below. We assume that in the transition from pre-malignant to malignant lesions, cells in a spatially well-structured population such as an epithelium are killed and/or extruded by an environmental stressor at a rate that is proportional to the inverse of their fitness, and replaced within the tissue via the division of a neighbouring cell. This assumption is represented in our chosen update rule. We suppose that it is this increased rate of cell turnover that leads to the accumulation of mutations, and eventually cancer. We refer to mutations as *advantageous, deleterious* or *neutral,* if they increase, decrease, or leave cell fitness unchanged.

The state of the system changes over time as a result of ‘death-birth’ events. At each point in time, each lattice site is defined by the presence of a cell with a specified ‘genotype’, given by the numbers of advantageous, neutral and deleterious mutations that it has accumulated. To implement the next death-birth event, we first choose a cell to die, at random, with a probability weighted by the inverse of each cell’s fitness. We define the fitness of a cell with *n_p_* advantageous, *n_n_* neutral, and *n_d_* deleterious mutations by 

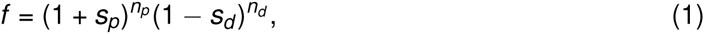
where the advantageous parameters *s_p_* and *s_d_* denote the relative fitness increase/decrease due to a advantageous/deleterious mutation. The chosen cell is removed from the lattice and one of the dead cell’s neighbours is chosen uniformly at random to divide into the vacated lattice site. The time at which this death-birth event occurs is given by a waiting time, chosen according to an exponential distribution with mean equal to the sum of all cell inverse fitnesses present on the lattice, as stipulated by the Gillespie algorithm [30].

Immediately following division, each daughter cell can independently accrue a mutation, with probability *μ*. If a mutation is accrued, it is labelled as advantageous, deleterious or neutral with equal probability 1/3. We note that neutral and deleterious mutations are not typically included in spatial Moran models of tumour evolution, as such mutations are unlikely to persist. However, over shorter timescales, their presence may have an effect on the dynamics of the system and hence the predictive power of any biomarkers considered. We emphasize that a cell that has accumulated mutations behaves the same as a wild-type cell in terms of mode of division and accumulation of mutations; the only difference between cells lies in their relative fitness, and hence the probability that they are chosen for removal as specified by the death-birth process.

We define the *time of clinical detection* of cancer to be the earliest time at which the proportion of cells with at least *N_m_* advantageous mutations exceeds a specified threshold *δ*. This reflects the time taken to reach a small, but clinically detectable, proportion of cancer cells that are capable of initiating and driving further tumour growth. In all simulations, we take *δ* = 0.05. We evaluate the correlation of a measurement of some property of the state of the lesion sampled at some time *T_b_* with the subsequent waiting time to cancer.

Measurements of the state of the lesion are performed by (i) taking a ‘biopsy’ from the lesion, and (ii) evaluating a putative ‘biomarker assay’ on the biopsy. Three different biopsy strategies are considered. First, we consider the whole lesion, in order to establish an upper bound on the prognostic power of each biomarker when using maximal information about the state of the system at a given time. Second, we sample a biopsy comprising a circular region of cells of radius *N_b_* lattice sites, whose centre is chosen uniformly at random such that the entire biopsy lies on the lattice; this represents the clinical procedure of core needle sampling. Third, we sample *N_s_* cells uniformly at random from the lattice; this represents washing or mechanical scraping of the lesion. In each case, we suppose that the biopsy constitutes a ‘snapshot’, and do not remove the sampled cells from the tissue. This simplifying assumption avoids the need to explicitly model the tissue response to wounding. The various biomarker assays evaluated are detailed below.

The definitions and values of all model parameters are summarized in Table 1. A MATLAB implementation of our model and simulated biopsy analysis is provided (see Text S1).

**Table 1.**
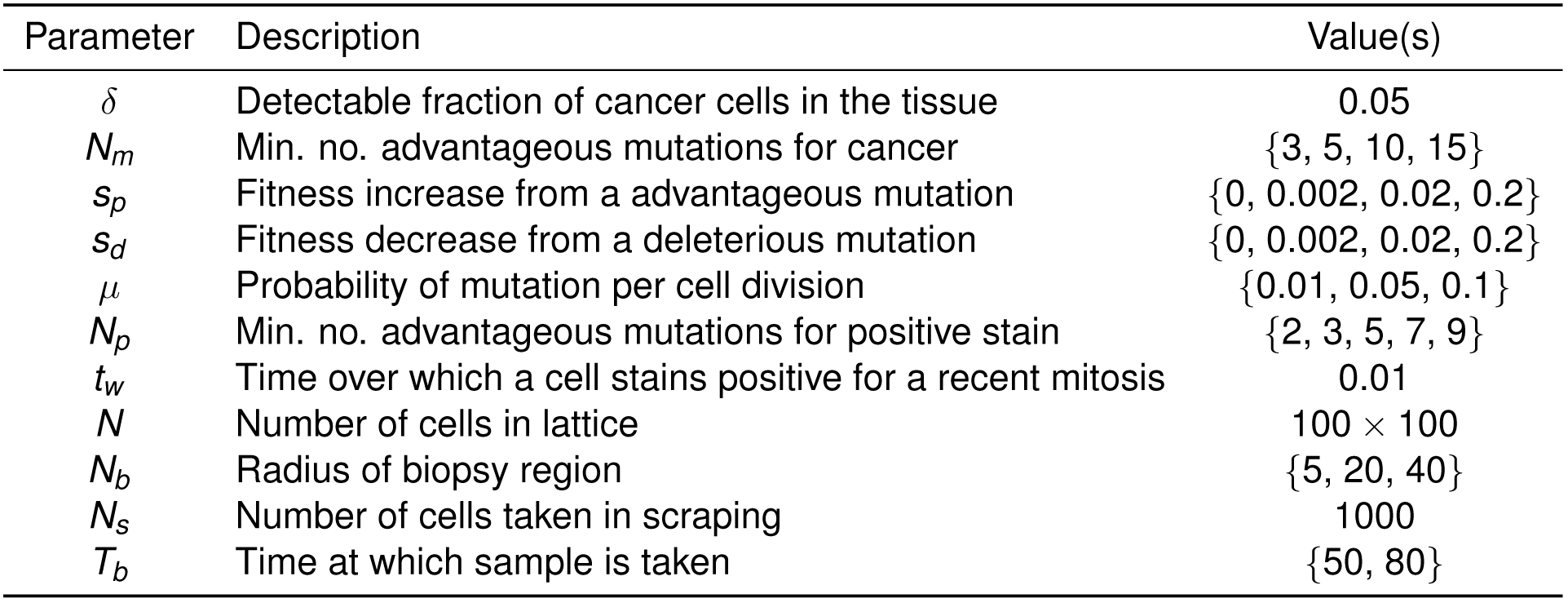
Parameter values used in the model.

### Classical biomarkers

**Proportion of cells with at least two advantageous mutations.** A commonly used class of biomarkers measure the proportion of cells in a biopsy staining positive for a given receptor. Examples include the estrogen receptor (ER), progesterone receptor (PR) and HER2/neu amplification staining commonly performed for malignancies of the breast [31, 32, 33]. Such assays are cost-effective and relatively simple to implement.

Here, we use the cutoff of a cell having acquired at least *N_p_* advantageous mutations to be representative of a cellular change that is observable in this manner. The measure is calculated simply by the number of cells having at least *N_p_* advantageous mutations, divided by the total number of cells sampled. We present results based on *N_p_* = 2 throughout the text, thus using the shorthand *N_p_* > 1 to refer to this biomarker. We discuss the robustness of these results to the chosen value of *N_p_* = 2 in the Results section.

**Mitotic proportion.** Proliferative cells are usually identified in tissue sections or cytology specimens using immunohistochemistry for cell-cycle associated proteins, foremost Ki-67 [34]. These proteins have a natural half-life over which a proliferative cell can be identified. To represent this measure in our computational model, we defined a time window *t_w_* over which a proliferative marker can be detected by staining. The mitotic proportion at a given time *t* is then defined as the number of cells that have undergone mitosis at least once in the time interval (*t* – *t_w_*, *t*], divided by the number of cells in the lattice, *N*.

### Measures of heterogeneity

**Shannon index.**The Shannon index *H* measures diversity among a population comprising different types [35]. For a population of *K* distinct types, each comprising a proportion *p_k_* of the population, the Shannon index is defined as

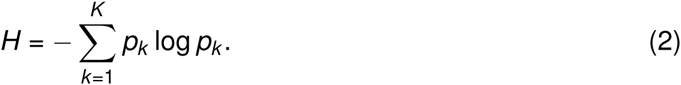

To calculate *H* we define *p_k_* such that each distinct triplet of advantageous, neutral and deleterious mutations is associated with a distinct clone within the model, and *p_k_* represents the proportion of cells in this clone.

**Gini-Simpson index.** Another established measure of diversity is the Simpson index [36]. To ensure that a higher value corresponds to greater diversity, we choose to use a transformation of this index called the Gini-Simpson index, *S*, which is defined as follows [37]. For a population of *K* distinct types, each comprising proportion *p_k_* of the population, we have

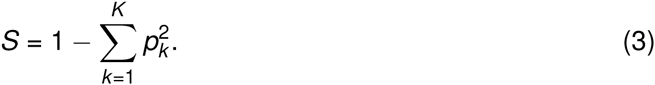

This index may be thought of as the probability that two randomly chosen members from the population are of different types. As the index increases towards the maximum of 1, the evenness of the distribution of the population over the various types becomes increasingly skewed toward one type.

**Moran’s *I*.** Moran’s *I* is a measure of global spatial autocorrelation which computes a weighted statistical average of the deviation between data points in a set, weighted by their spatial distance [38]. Moran’s *I* takes values in [−1, 1]. For a given set of values {*X*_1_... *X_N_*}, with mean 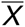, and spatial weight matrix 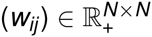, Moran’s *I* is defined as

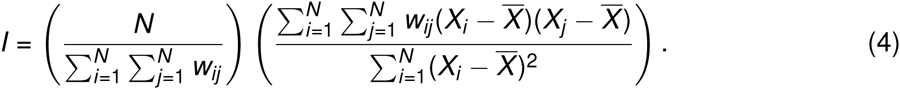

We take *X_i_* to be the sum of the numbers of advantageous, neutral and deleterious mutations accumulated by the cell at lattice site *i*. Clinically, *X_i_* may be thought of as a binary variable (0 or 1) indicating whether a given cell bears some detectable abnormality (for instance a particular number of advantageous mutations). The spatial weight matrix (*w_ij_*) can be specified in several ways; here, we define 

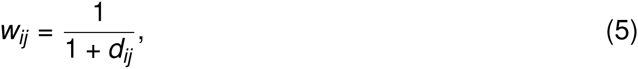
where *d_ij_* is the Euclidean distance between the lattice sites indexed by *i*, *j* ∈ {1,..., *N*}. With this functional form, neighbouring points that are closer together are weighted more heavily, thus contributing more to the measure.

**Geary’s** *C.* Geary’s *C,* like Moran’s *I*, is a global measure of spatial autocorrelation. Geary’s *C* takes values in [0, 2], with higher values indicating less spatial autocorrelation, and lower values indicating a greater degree of spatial autocorrelation [39]. While Moran’s *I* is a more global measurement and sensitive to extreme observations, Geary’s *C* is more sensitive to differences in local neighbourhoods. For a given set of values {*X*_1_... *X_N_*}, with mean 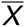, and a given spatial weight matrix 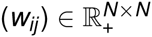, Geary’s *C* is defined as

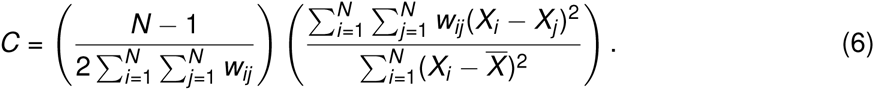

Here, our definitions of *X_i_* and (*w_ij_*) follow those given for Moran’s *I.*

**Index of positive proliferation (IPP).** We next define a novel measure, termed the index of positive proliferation (IPP), that is a spatially weighted average of the location of mitotic cells and the number of advantageous mutations accrued by nearby cells. The biological motivation for this measure is to detect the recent clonal expansions of advantageous mutants, in order to quantify evidence of recent progression towards cancer: we might expect the concentration of proliferation in regions of high numbers of advantageous mutations to correlate with a poor prognosis.

We define the IPP as follows. Consider a population of *N* cells, with the individual cells labelled as *X*_1_,..., *X_N_*. Suppose that a subset of these cells, *Y*_1_,..., *Y_Q_*, are proliferating at a given time. We define *cellular contributions f*_1_,..., *f_M_* ∈ ℝ^+^ as values such that a higher contribution corresponds to a cellular state genetically closer to that of cancer. Clinically, these cellular contributions correspond to cells that are genotypically closer to the end state of cancer, and represent either cutoff points that may be detected by gene sequencing, or immunohistochemical changes. For each cell *i* with *m_i_,* advantageous mutations, we define the cellular contribution *f_i_* as

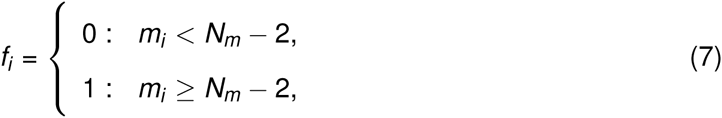

Thus, for a given spatial weight matrix (*w_ij_*) ∈ ℝ*^N^*^×^*^N^* we define the IPP as

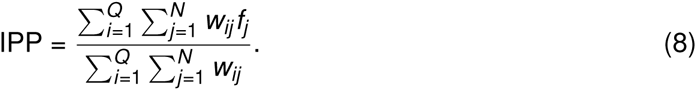

We define the weights *w_ij_* as in equation (5), where *d_ij_* is the Euclidean distance between cell *X_i_* and proliferating cell *Y_j_*, such that *X_j_* ≠ *Y_j_*. In the case that *X_i_* = *Y_j_*, we take *W_ij_* = 0.

Since time steps correspond to ‘generations’ in our mathematical model, we store the locations of the most recent cell divisions within a given time window *t_w_*, and regard these locations as locations that would stain positive for mitotic activity (e.g. via Ki-67 staining), to model the non-instantaneous process of detecting active cell division.

**Index of non-negative proliferation (INP).** To model the case where it may not be feasible to observe an accumulation of advantageous mutations only, in the sense that mutations accumulated may be neutral as well, we define an additional measure termed the index of non-negative proliferation (INP). This measure is defined analogously to the IPP, but with the cellular contributions chosen such that 

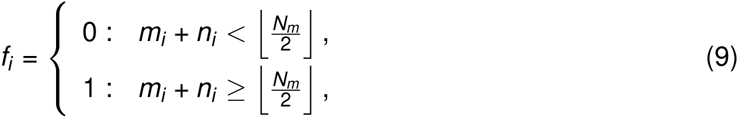
where ⌊·⌋ denotes the integer part. Here, the sum of the number of advantageous and neutral mutations is considered to be the observable quantity, simulating a situation in which the observable information encapsulates and may skew the perception of the true genotypic state of the system. The cutoff value of *N_m_*/2 was chosen in an *ad hoc* manner based on preliminary simulations; we note that refinement of this parameter may be necessary for effective use of the INP in future studies.

### Statistical methods

The statistical association (correlation with the waiting time to detectable cancer) of each sampling strategy and putative biomarker assay was evaluated using Kaplan-Meier curves and univariate Cox Proportional Hazards models as implemented in the *R* statistical computing language. For all presented p-values, the significance cutoff is taken as 0.05.

Data used for the Cox regression model were all generated by the stochastic simulations of the computational model. That is, the event times were defined as the simulation times at which 5% of the cells of the lattice were defined as cancerous, and the predictors of this time were taken to be the biomarker index values computed at an earlier simulation time. The cohort size is therefore the number of such simulations which were carried out, which was 10^3^. There was no censoring required, as all simulations were run to completion of endpoint as defined previously.

## Results

We consider a spatial model of the evolution of malignancy in a precancerous lesion. In our model, cells occupy a two-dimensional lattice of size *N*. Time is treated as a continuous variable in the model, but is simulated as a succession of discrete time steps, where the length of each time step is a function of the overall fitness of the population and a stochastic factor, as per the Gillespie Algorithm [30]. At each time step, a cell is chosen at random to die and is removed from the lattice with a probability that is inversely proportional to its *cellular fitness,* a positive real number that is initially equal to 1 for non-mutated cells and may be altered by mutation. When a cell dies, one of its neighbours is then chosen uniformly at random to divide, with one of the daughter cells occupying the free lattice site and each daughter cell independently acquiring a new mutation with probability *μ*. We refer to mutations as *advantageous, deleterious* or *neutral,* according to whether they increase, decrease, or leave fitness unchanged, with each type of mutation assumed to be equally likely.

Starting from a lattice occupied entirely by non-mutant cells, we consider the *outcome* of each simulation to be the time taken for the proportion of cells with at least *N_m_* advantageous mutations to exceed a threshold *δ*. This waiting time is defined as the time to clinically detectable lesions. We choose a value of *δ* corresponding to a proportion of cancer cells that is sufficiently large to be clinically detectable, and to initiate subsequent rapid growth.

A representative snapshot of a model simulation is shown in Fig. 1A. To simulate clinical sampling, at a predetermined time *T_b_* we take a virtual *biopsy* from the lesion (Fig. 1B), from which we compute various biomarkers and assess their prognostic value in determining the waiting time to cancer (see *Methods*). The model exhibits successive clonal sweeps of mutations (Fig. 1C).

**Figure 1.**
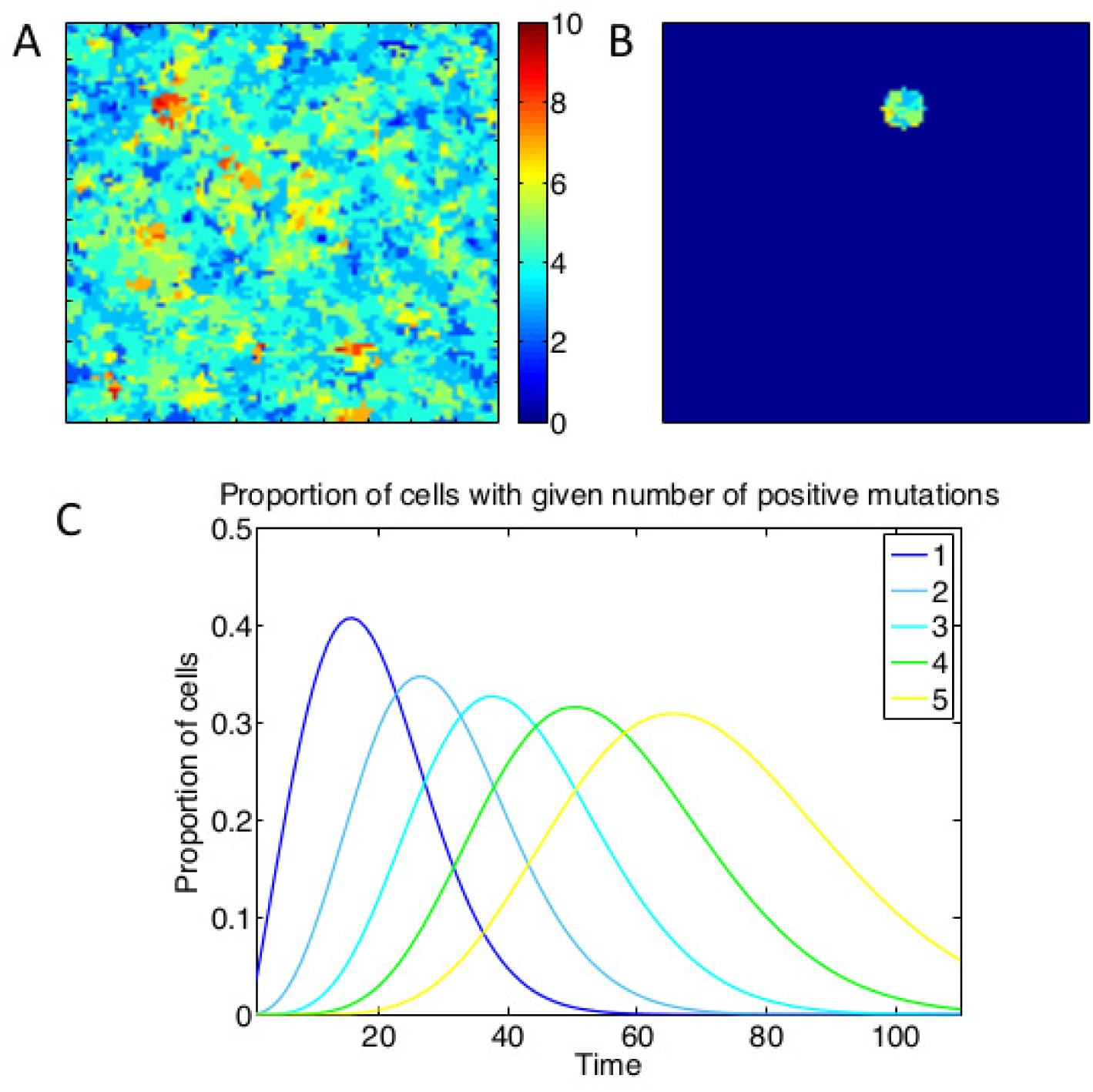
Depiction of the spatial simulation, a virtual biopsy, and the successive clonal sweeps. A: Heat map of the lattice at a given point in time, with different colours representing different numbers of positive mutations of the cells at those points. B: Depiction of the lattice subset involved in a virtual biopsy. C: Time evolution of the proportions of cells with different numbers of positive mutations, showing successive clonal sweeps. Results are averaged from 200 simulations with parameter values *N_m_* = 10, *s_p_* = *s_d_* = 0.2 for five such genotypes (for figure clarity).

### Assessment of candidate biomarkers and tissue sampling schemes

**Counting driver mutations.** We considered the correlation between the proportion of cells bearing at least *N_p_* advantageous mutations (so-called driver mutations) and the waiting time to cancer. The closer the cutoff *N_p_* is to the number of mutations required for cancer, *N_m_*, the more correlated this measure became with the waiting time to cancer (Table S1). These results confirm the intuition that it is easier to predict the occurrence of a cancer at a time point close to when the cancer will occur (e.g. at the ‘end’ of the evolutionary process, when only a few additional driver mutations are required) than early in the cancer’s development, when many additional mutations are required.

**Small needle biopsies.** We computed the prognostic value of various candidate biomarker ‘assays’ performed on a single biopsy of radius *N_b_* = 20 taken at time *T_b_* post simulation initiation. Neither the proportion of cells with at least one advantageous mutation nor the proliferative fraction were significant predictors of prognosis (Table 2). In contrast, measures of clonal diversity (Shannon and Gini-Simpson index) were both highly significant predictors of prognosis (*p* < 10^−4^ in both cases). Of the spatial autocorrelation measures, Moran’s *I* (*p* = 0.02) but not Geary’s *C* (*p* = 0.29) had prognostic value.

**Table 2.** Summary of Cox proportional hazards models for various putative biomarker schemes, for different tissue sampling schemes. Hazard ratios (HR75) are computed at time *t* = 75 for the case *N_m_* = 10, *s_p_* = *s_d_* = 0.2, and *μ* = 0.1. Statistically significant values are in bold. ‘Unit change’ denotes the change in the value of each putative biomarker that increases the associated hazard ratio by the reported factor.

| | | Unit change | HR75 | 95% CI | $p$ |
| --- | --- | --- | --- | --- | --- |
| Whole lesion | $n_p > 1$ proportion | 0.05 | 0 | $(0, \infty)$ | 0.29 |
| | Mitotic proportion | 0.01 | **0.31** | $(0.12, 0.81)$ | 0.02 |
| | Shannon index | 0.1 | **2** | $(1.8, 2.2)$ | $< 10^{-4}$ |
| | Gini-Simpson index | 0.01 | **5.5** | $(4.2, 7.2)$ | $< 10^{-4}$ |
| | Moran's $I$ | 0.05 | **3.2** | $(2, 5.3)$ | $< 10^{-4}$ |
| | Geary's $C$ | 0.01 | 0.98 | $(0.92, 1)$ | 0.43 |
| | IPP | 0.01 | **2** | $(1.9, 2.1)$ | $< 10^{-4}$ |
| | INP | 0.01 | **1.3** | $(1.2, 1.4)$ | $< 10^{-4}$ |
| Biopsy | $n_p > 1$ proportion | 0.05 | 0.95 | $(0.86, 1.1)$ | 0.35 |
| | Mitotic proportion | 0.01 | 0.81 | $(0.58, 1.1)$ | 0.22 |
| | Shannon index | 0.2 | **1.3** | $(1.1, 1.4)$ | $< 10^{-4}$ |
| | Gini-Simpson index | 0.01 | **2.3** | $(1.5, 3.5)$ | $< 10^{-4}$ |
| | Moran's $I$ | 0.1 | **1.4** | $(1.1, 1.9)$ | 0.02 |
| | Geary's $C$ | 0.1 | 0.95 | $(0.87, 1)$ | 0.29 |
| | IPP | 0.01 | **1.1** | $(1, 1.1)$ | $< 10^{-4}$ |
| | INP | 0.1 | 1 | $(0.95, 1.1)$ | 0.57 |
| Scraping | $n_p > 1$ proportion | 0.05 | $< 10^{-6}$ | $(0, 0.0001)$ | 0.01 |
| | Mitotic proportion | 0.01 | 1.2 | $(0.88, 1.7)$ | 0.23 |
| | Shannon index | 0.05 | **1.3** | $(1.3, 1.4)$ | $< 10^{-4}$ |
| | Gini-Simpson index | 0.01 | **3.5** | $(2.8, 4.4)$ | $< 10^{-4}$ |

**Random sampling.** Random sampling of cells from the lesion represents a tissue collection method such as an endoscopic brush or a cellular wash. We took a random sample of 10^3^ cells, corresponding to 10% of the total lesion. As for small biopsy sampling, the proportion of mitotic cells within the sample was a poor prognostic marker (*p* = 0.23), but interestingly the proportion of cells with more than one advantageous mutation became a significant predictor (*p* = 0.01). This may be due to the fact that within a sparse sample of the lesion, the number of mutant cells is a proxy for active on-going evolution: either via the large scale clonal expansion of a single clone, or multiple foci of independent clones. Increased clonal diversity remained a highly significant predictor of a short waiting time to cancer (*p* < 10^−4^ for both the Shannon and Gini-Simpson indices; Fig. 2).

**Figure 2.**
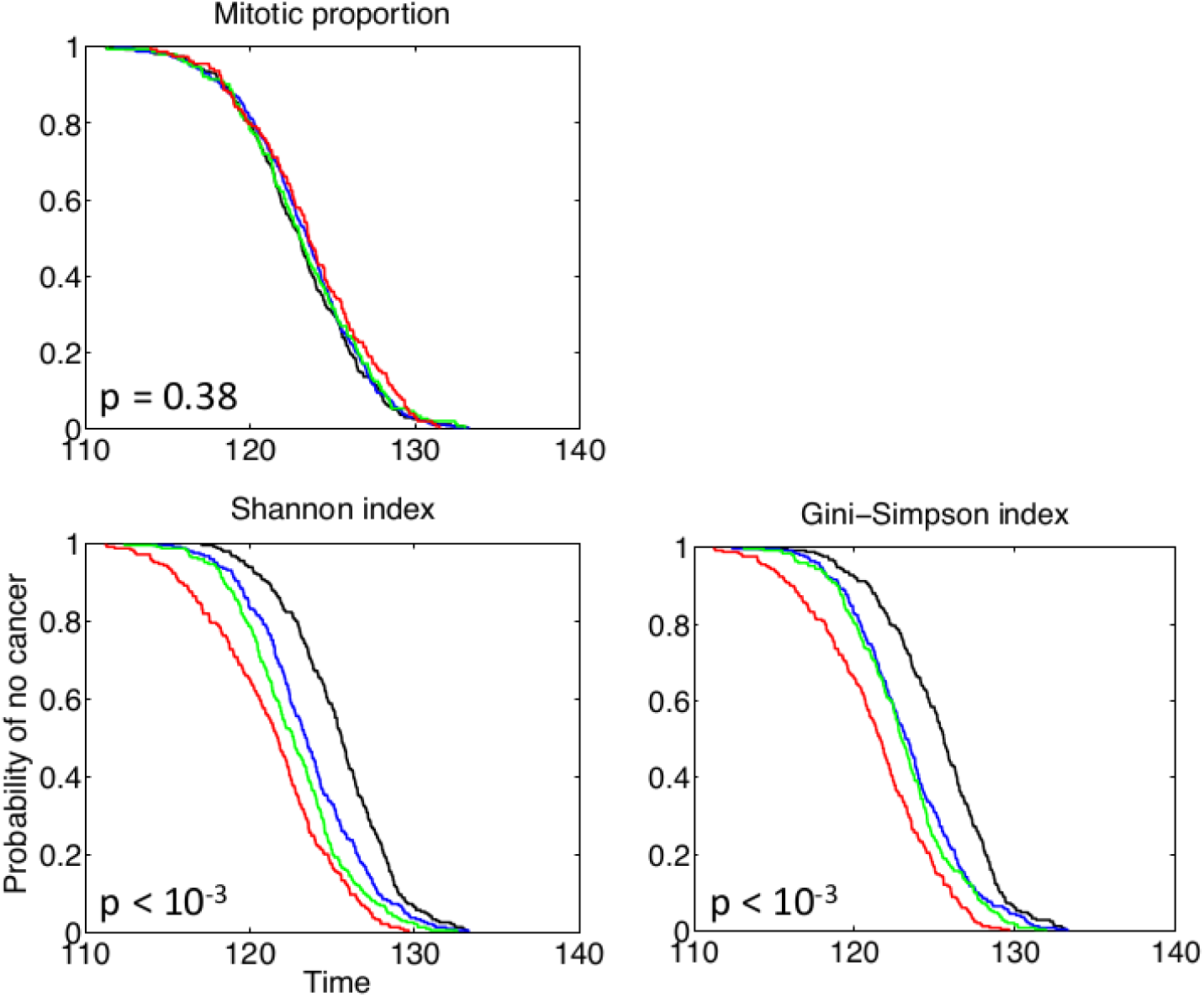
Prognostic value of random tissue sampling. A random sample of *N_s_* = 10^3^ (10% of the lesion) cells was sampled at time *T_b_* = 80 and the prognostic value of the mitotic proportion (A), Shannon index (B) and Gini-Simpson index (C) on this sample was considered. Kaplan-Meier curves are plotted for each putative biomarker assessed, and in case, the values across the simulations were separated into upper (red), upper middle (green), lower middle (blue) and lower (black) quartiles. Only biomarkers that did not require spatial information could be computed for this tissue sampling method. *P*-values are for the generalized log-rank test.

**Whole lesion sampling.** In the case of whole-lesion sampling, all information on the current state of the virtual tumour is available in the biomarker assay, and hence we expected to see maximum predictive value of our putative biomarkers. In this case, the proportion of cells with at least one advantageous mutation remained a poor prognosticator (*p* = 0.29), whereas the proportion of proliferative cells became a significant predictor (*p* = 0.02) (see Table 2).

The clonal diversity measures remained highly significant prognosticators (*p* < 10^−4^ in both cases), underlining their robustness as prognostic measures. Higher clonal diversity was associated with faster progression to cancer (Fig. 3). The prognostic value of the spatial autocorrelation measure Moran’s *I* was significantly improved when the whole grid was sampled (*p* < 10^−4^), but Geary’s *C* remained non-correlated.

**Figure 3.**
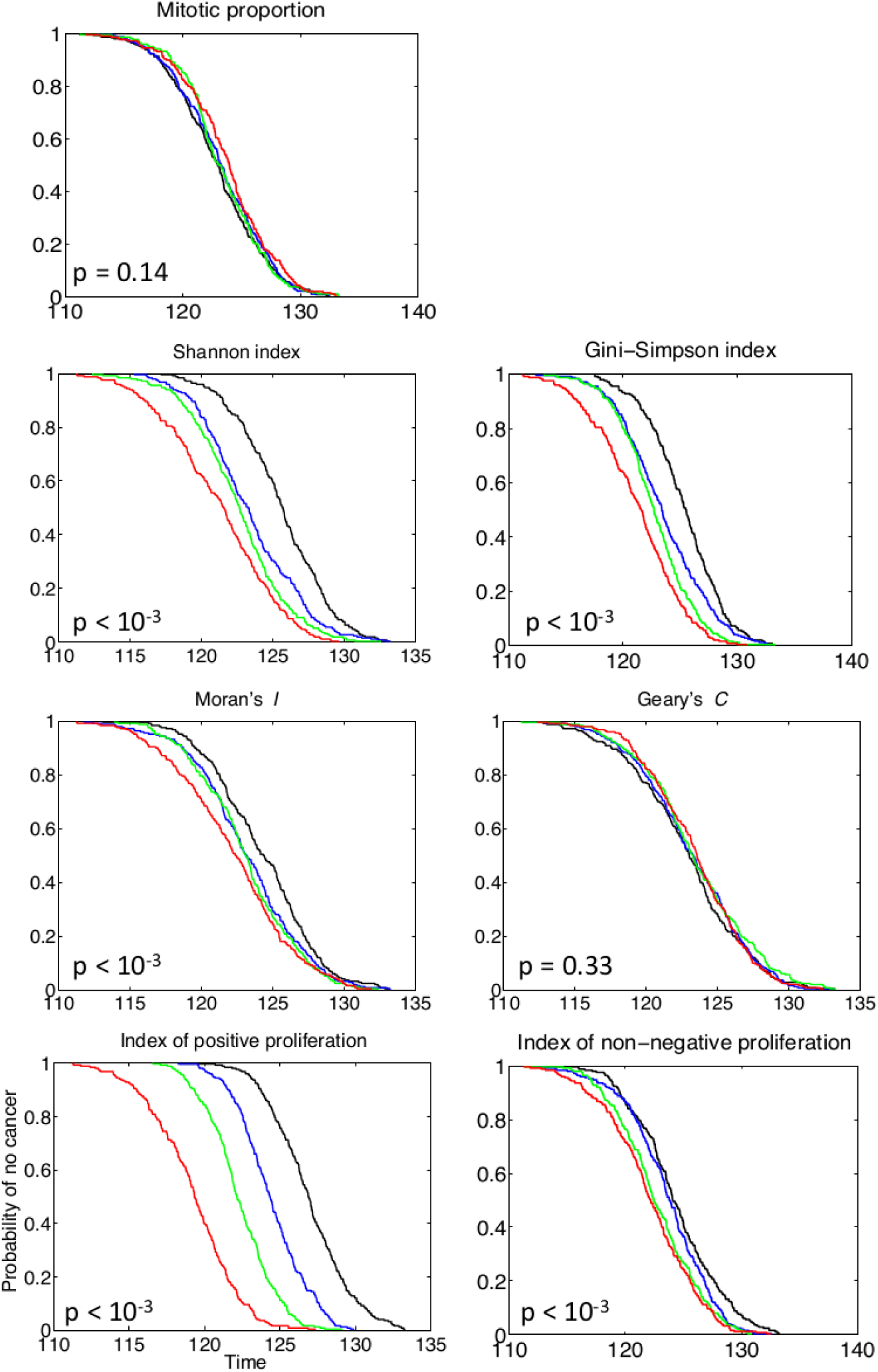
Sampling the whole lesion improves the prognostic value. The prognostic value of sampling the whole lattice at time *T_b_* = 80 was assessed. Kaplan-Meier curves are plotted for each putative biomarker assessed for biomarker values across the simulations were separated into upper (red), upper middle (green), lower middle (blue) and lower (black) quartiles. *P*-values are for the generalized log-rank test.

Together these data highlight the high prognostic value of diversity measures, and their robustness to the details of tissue sampling method used.

### Novel prognostic measures

We next sought to determine whether novel statistics calculated on the state of the lesion could provide additional prognostic value. We defined two new statistics, the *index of positive proliferation* (IPP) and the *index of non-negative proliferation* (INP), which describe the spatial autocorrelation between proliferating cells with advantageous mutations, or proliferating cells with non-deleterious mutations, respectively (see *Methods*). Since these statistics tie together measures of both the mutation burden and proliferative index, we consider them to be measures of the degree of ‘evolutionary activity’.

In both small biopsy samples and whole-lesion analysis, the IPP was a highly prognostic statistic (Table 2), with larger values of the statistic accurately predicting shorter times to cancer (Fig. 3). The INP was prognostic on whole-lesion analysis (Fig. 3), but not on targeted biopsies (Table 2). The difference in the prognostic value between IPP and INP is suggestive of the particular importance of assaying ‘distance’ travelled along the evolutionary trajectory towards cancer: the IPP is sensitive to this distance as it only measures advantageous mutations, whereas the INP is potentially confounded by non-adaptive mutations. The inherent issues associated with the identification of advantageous mutations consequently potentially limit the utility of these novel measures.

To assess the predictability of each putative biomarker [40] we calculated the area under the receiver operating characteristic (ROC) curves (Fig. 4) as a function of the censoring time [41]. ROC curves are the curves defined by the sensitivity and specificity of each index value as it predicts end time to cancer, where a positive end time is a time past a certain pre-defined simulation time, and the cutoff for the index value that defines whether the index predicts if that end time is early or late, is continuously varied. The area under these curves is 1 in the case of an index value that is perfectly predictive of an end time, and 0.5 for random guessing as to whether the index predicts if the end time is early or late. These curves show the IPP measure has the best predictive value of all measures considered, and that the Shannon and Gini-Simpson diversity indices also have strong predictive value. The lack of predictive value derived from the mitotic proportion, Geary’s *C* and proportion of mutant cells was also confirmed.

**Figure 4.**
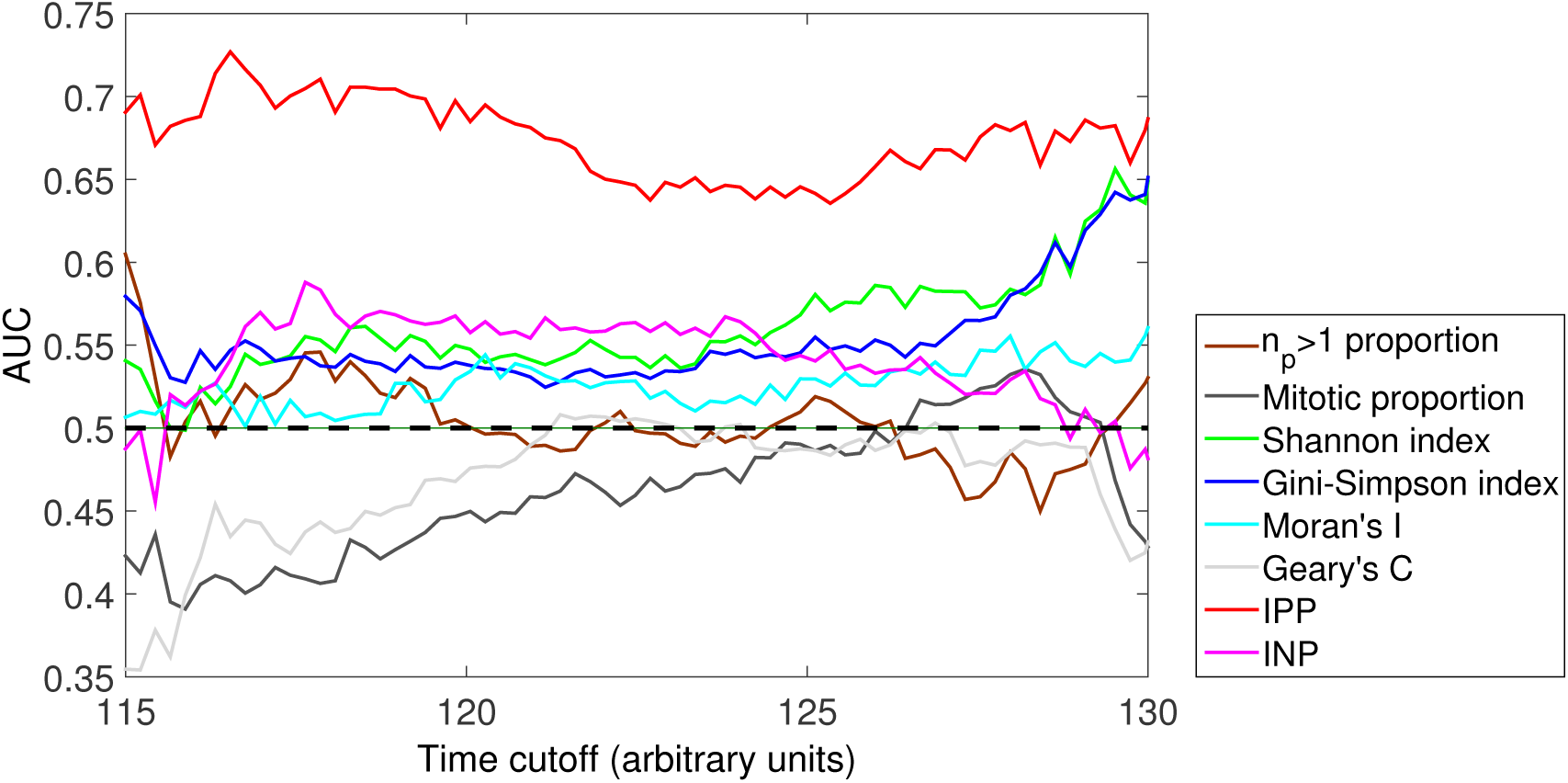
Areas under ROC curves for putative biomarkers. The prognostic value of sampling a circular biopsy at time *T_b_* = 75 was assessed by considering the area under the curve (AUC) of receiver-operator characteristic (ROC) curves as a function of censoring time. This analysis confirmed the time-invariant predictive value of the IPP (red line) and clonal diversity measures (blue and green lines), and lack of predictive value derived from the mitotic proportion (black line) and proportion of cells bearing at least one abnormality (brown line). The worse-than-random performance of the proliferation and Geary’s *C* measurs at short censoring times is likely to be attributable to the stochasticity inherent in cancer development within the model: early clonal expansions do not necessarily signify later cancer risk. Results from 1000 simulations for each sampling scheme, with parameter values *N_m_* = 10, *s_p_* = *s_d_* = 0.2, *μ* = 0.1, *N_b_* = 20 and *N_s_* = 10^3^. For comparison, the black dotted line denotes an AUC=0.5 (which would would be achieved by a random predictor).

### Early versus late biopsy

Effective screening for cancer risk requires predicting cancer risk long before the cancer develops. We next considered how the timing of a biopsy affects its prognostic value by investigating how the correlation coefficient between each biomarker and the subsequent waiting time to cancer varies with the time at which the biopsy is taken.

As expected, we found that biopsies collected later in the lesion’s evolution (e.g. closer to the time of cancer development) generally had more statistical association than biopsies collected earlier, and this was true irrespective of the tissue sampling method used (Fig. 5A-C). Sampling early in the lesion’s evolution (e.g. near to the start of the simulations) had poor correlation irrespective of the putative biomarker assay used, reflecting the fact that very few mutations had accumulated in the lesion at short times. Sampling at intermediate times showed dramatic improvements in the prognostic value of the diversity indices and IPP measure, whereas samples taken at a variety of long times had approximately equal prognostic value or showed slight declines relative to intermediate times. At intermediate and long times, the IPP was the best performing prognostic measure. The mitotic proportion and proportion of cells with at least one advantageous mutation were consistently poor predictors across the entire time course.

**Figure 5.**
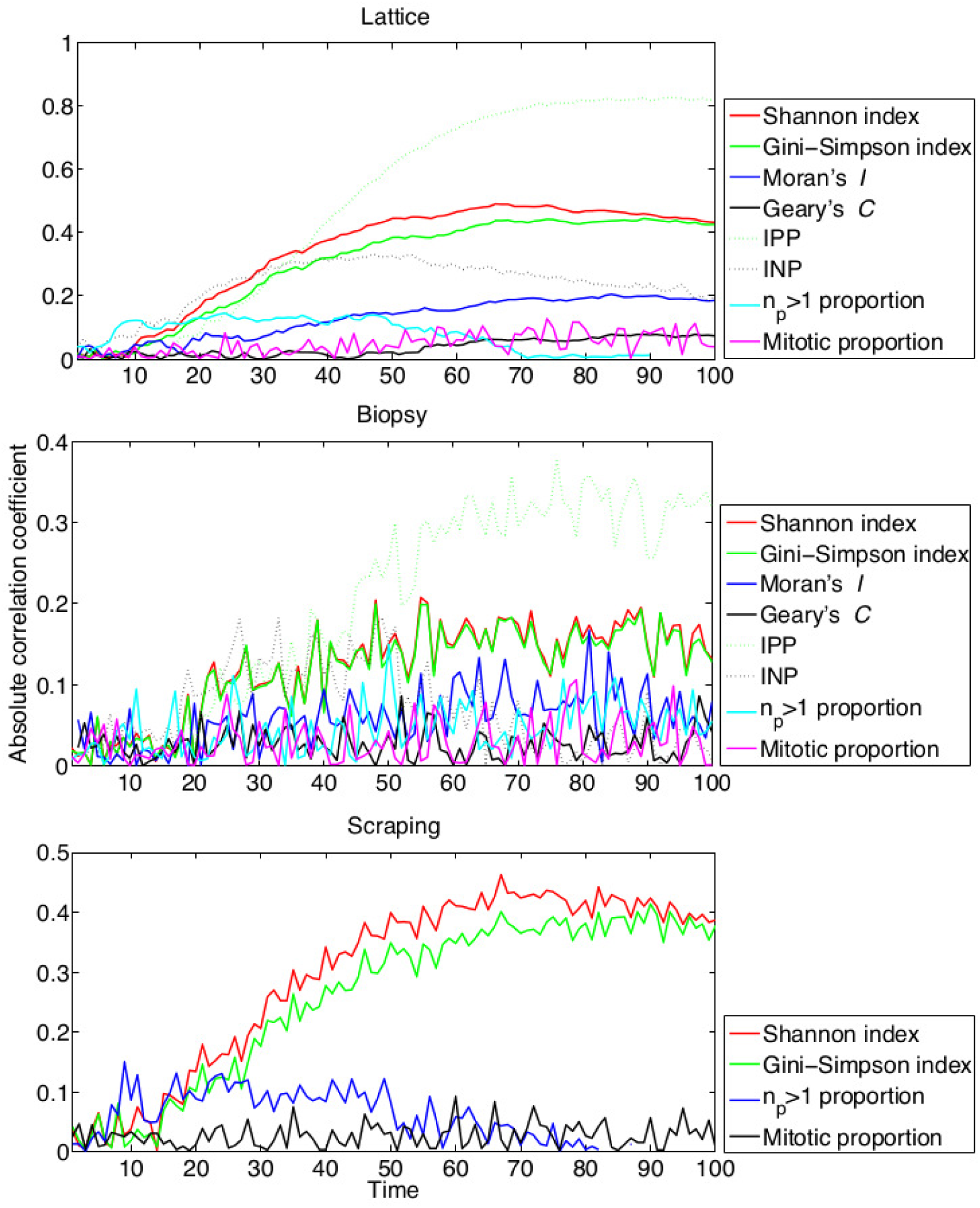
Prognostic value of early versus late biopsies. For a range of sampling times *T_b_*, the virtual tissue was biopsied and the correlation between putative biomarker values and the waiting time to cancer was computed. Results from 1000 simulations for each sampling scheme, with *N_m_* = 10, *s_p_* = *s_d_* = 0.2, *μ* = 0.1, *N_b_* = 20 and *N_s_* = 10^3^.

The effect of taking a small biopsy, as opposed to sampling the whole lesion, was to both significantly reduce the prognostic value of all putative biomarker measures, and introduce ‘noise’ into their prognostic values (Fig. 5B). Importantly, we observed that in spite of this noise, the correlation coefficients for the clonal heterogeneity and IPP measures were consistently high compared to the other measures, indicating their robustness as prognostic markers. Biopsy sampling significantly reduced the prognostic value of Moran’s *I* compared to whole-lesion sampling, indicating how this measure is particularly confounded by tissue sampling.

On random samples (analogous to endoscopic brushings or washings), the Shannon and Gini-Simpson indices showed good correlations with the waiting time to cancer. These diversity measures were more correlated for random samples than for circular biopsies, despite each sample constituting similar numbers of cells (10% and 12% of the lesion, respectively). This result may reflect the fact that a biopsy can potentially miss a ‘dangerous’ clone, whereas a random sampling method is likely to obtain cells from all sizeable clones within the lesion.

Together, these data indicate that larger samples usually provide more prognostic value than smaller samples, and that very ‘early’ tissue samples are unlikely to contain significant prognostic information. They also highlight again that the prognostic value of diversity measures is particularly robust to the details of tissue sampling.

### Longitudinally collected biopsies

We next examined whether combining information from serial biopsies, taken at two different time points (*t*_1_ and *t*_2_; both strictly before cancer occurrence), provided more prognostic information than a biopsy from a single time point. To do this, we evaluated the average of the values of each biomarker at the times *t*_1_ and *t*_2_, and the correlation between this average and the waiting time to cancer. We then compared this correlation with that of the biomarker value at time *t*_2_ alone. These results are shown in Fig. 6, where the *x* and *y* axes indicate the time of the first and second biopsies, and the colours indicate the difference between the correlation coefficient for the *average* of the individual biomarker values at each time points and the correlation coefficient for the biomarker value at the second time point alone.

**Figure 6.**
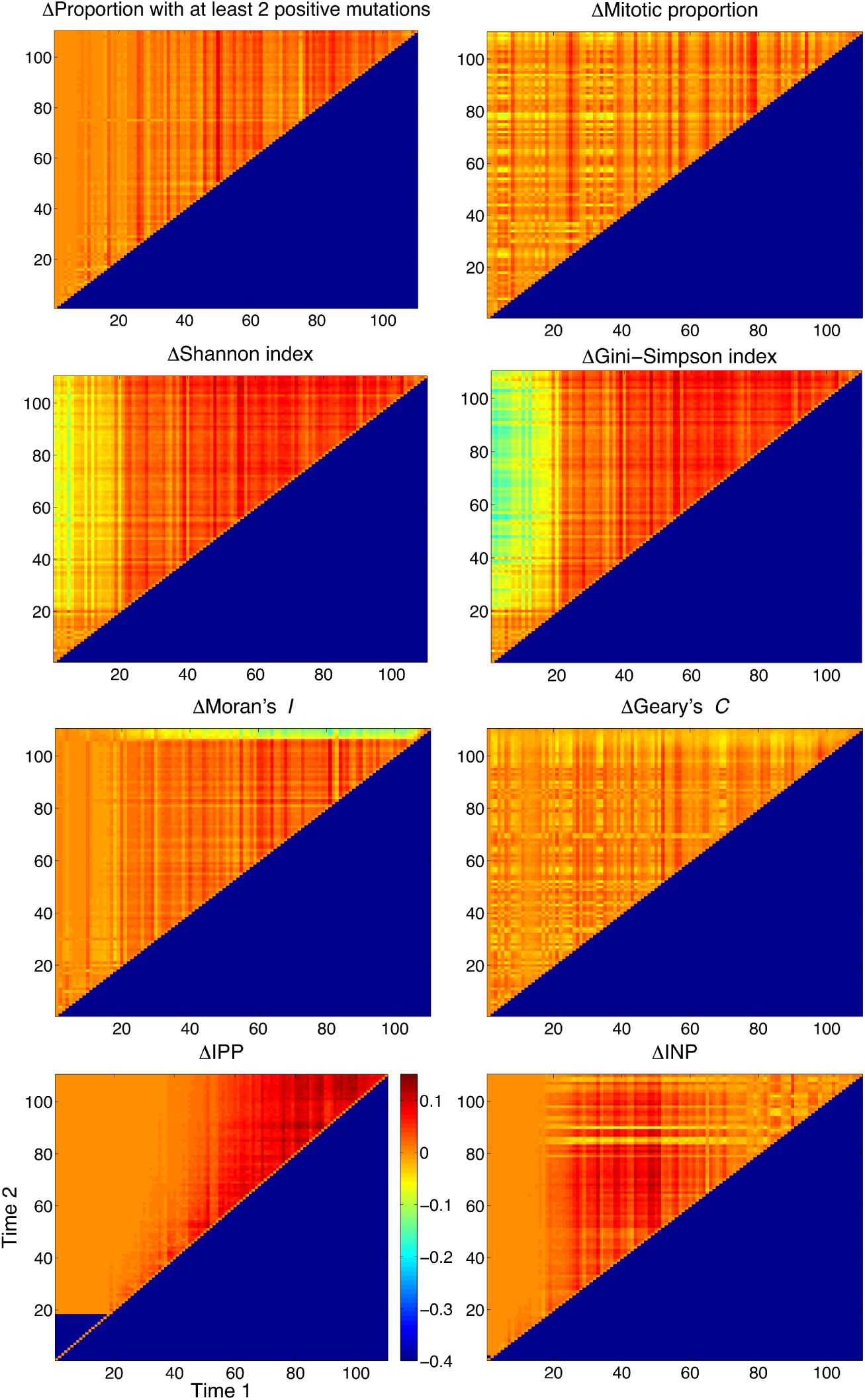
Serial biopsies provide slightly increased additional prognostic information. Heat maps depicting the relative value of taking serial biopsies at different time points. Positive values (warm colours) indicate that prognostic value was improved by taking the average of biomarker value from both time-points; negative values (cool colours) indicate that more information was available at the second time point alone than from the averaged time points. Results from 1000 simulations for each pair of time points, with *N_m_* = 10, *s_p_* = *s_d_* = 0.2, *μ* = 0.1 and *N_b_* = 20.

Including information from an early biopsy in this manner provided slight additional prognostic value over-and-above the information available in the later biopsy (Fig. 6; approximately a 0.1 increase in the correlation was observed). In contrast, When information was combined by taking the difference in biomarker values between the biopsies collected at two different time points, the value from the later biopsy was generally more prognostic, and importantly was more prognostic than a measure which combined information both the early and late biopsies. Interestingly, the prognostic value of the Shannon and Gini-Simpson indices was reduced when considering the difference in biomarker values between two time points (Fig. S4). When we instead compared the maximal value of the biomarker across the two time points to its value at the later time point, we found similar results to the average case, but with smaller increases in correlation at later times; this was particularly the case for the Shannon, Gini-Simpson, IPP and INP indices (Fig. S5).

**Multiple biopsies at the same time point**

A consequence of intra-tumour heterogeneity is that a single biopsy may fail to sample an important clone [42] and so cause an incorrect prognosis assignment. To address this issue, we studied how the prognostic value of each putative biomarker was improved by taking additional biopsies at the same time (*T_b_* = 50). For simplicity, after each virtual biopsy the sampled tissue was perfectly replaced in order to avoid the complexities associated with modelling local wound healing and tissue recovery. Further, while we did not strictly preclude biopsies from overlapping, the degree of overlap between biopsies is typically minimal because of the relatively small numbers of biopsies and small sizes of biopsy considered.

Assaying from more biopsies generally improved prognostic value, but with diminishing returns for each additional biopsy (Fig. 7). For all but one of the putative biomarkers, the maximum prognostic value was achieved by taking the average biomarker value across all biopsies, whereas measures of the spread of values (the variance or range) were generally poor prognosticators. Interestingly, the maximum prognostic value for the proportion of cells with at least two advantageous mutations was achieved by taking the *minimum* value across all biopsies; this could be because the minimum value is particularly sensitive to biopsies that contain non-progressed cells. Together these data imply that taking more biopsies and averaging the biomarker signal across biopsies provides additional prognostic information.

**Figure 7.**
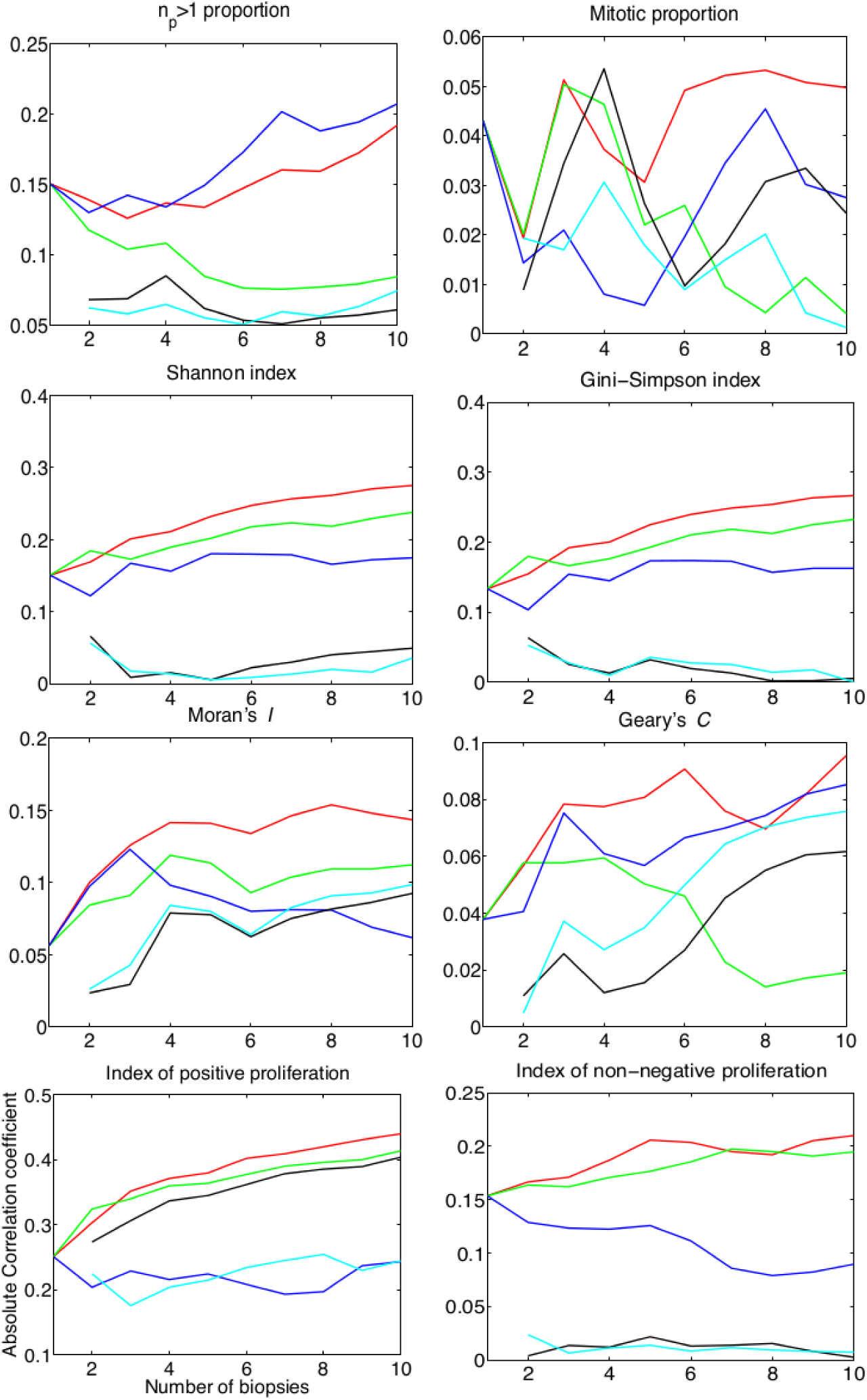
Additional biopsies at the same time point improves prognostication with diminishing returns. Graphs show the relationship between the correlation coefficient (between each biomarker value and waiting time to cancer) and the number of biopsies collected at time *T_b_* = 50. Lines denote different measures based on the multiple biopsies: average biomarker value across biopsies (red); maximum value (green); minimum value (blue); difference between maximum and minimum values (black); and variance in values (cyan).

### Robustness of results to choice of model

To assess the robustness of results to our model assumptions, we investigated the impact of parameter values and update rules on the statistical association of each biomarker. We observed the same qualitative behaviour, such as diversity measures outperforming the proliferative fraction in degree of correlation, irrespective of the choice of parameter values or update rule used (see Figs S1–S3, Tables S1–S17 and Text S1 for details).

To briefly summarize these results: (i) lower mutation rates decreased the correlation of each marker with the waiting time to cancer, because of the increased stochasticity in the model introduced by a lower mutation rate; (ii) smaller biopsies were in general less prognostic; (iii) the number of mutations required for cancer did not qualitatively change the predictions of the model; (iv) the fitness advantage and disadvantage caused by new mutations, and the relative likelihood of each of the various mutation types, did not qualitatively alter the prognostic value of the biomarkers, although diversity measures were most prognostic for the case where there were many strongly advantageous mutations; and (v) the closer the value of the threshold *N_p_* was to the number of mutations required for cancer, the more correlated the proportion of cells with *n_p_* ≥ *N_p_* became with the waiting time to cancer.

We also analyzed the sensitivity of model results to variations in the update rules of the system. We tested the biomarkers in a birth-driven system as opposed to a death-driven system, and found that again, that the diversity and IPP measures remained the biomarkers most significantly associated with the waiting time to cancer. Further, we investigated the effect of decoupling mutations and cell division. While these changes to the model did alter the specific predictive values of the various indices (summarized in Table S14), the general pattern of statistical association was not altered. Moreover, even in this scenario, the IPP performed well.

## Discussion

In this work we have developed a simple computational model of cancer development within premalignant disease and used the model to evaluate the prognostic value of a range of different putative biomarker measurements and tissue sampling schemes. Our results show that simply counting the proportion of cells bearing multiple advantageous mutations (proportion of cells with ‘driver’ mutations) or the proportion of proliferating cells were universally poor predictors of the waiting time to cancer, whereas measures of clonal diversity were highly correlated with the time to cancer and were robust to the choice of tissue sampling scheme. Further, we evaluated a range of different tissue sampling schemes (single biopsy, multiple biopsies in space or time, or random sampling of a lesion). We found that random sampling (such as via an endoscopic brush) provided more consistent prognostic value than a single biopsy, likely because a single (randomly targeted) biopsy is liable to miss localised but ‘important’ clones. Prognostication was improved by taking multiple biopsies, but with diminishing returns for each additional biopsy taken. Together these data provide a rationale for the empirical evaluation of different tissue sampling schemes.

Averaging biomarker scores from two different time points did improve the predictive value of our putative biomarkers; however, the difference in each putative biomarker’s values between time points was less predictive than its value at the later time points. This result was somewhat counter to our initial intuition that taking longitudinal biopsies would accurately track the ‘evolutionary trajectory’ of the lesion and hence dramatically improve prognostication. This result illustrates how our *in silico* approach can challenge intuition and, in so doing, provide novel insights into biomarker development.

We developed a new statistic, termed the index of positive proliferation (IPP), that proved to be a highly prognostic measure. The IPP is a measure of the average distance to a proliferating cell that has acquired advantageous mutations. It thus combines both genetic (or phenotypic) information with spatial (cell position) and dynamic (proliferation) information. This integration of multiple different sources of information may account for the prognostic value of the biomarker in our model. Empirical measurement of the IPP would be feasible if, for example, the number of driver mutations accumulated by a cell could be quantified concomitantly with a proliferative marker. Developments in *in situ* genotyping methods might facilitate such an approach in the near future. Irrespective of the immediate feasibility of such a measure, our development and testing of the IPP statistic within our computational model illustrates how *in silico* approaches provide a powerful means to rapidly explore new potential biomarker assays.

Our computational model of cancer evolution is clearly a highly simplified description of reality. For example, we modelled a simple two-dimensional sheet of epithelial cells and neglected the important influence, and indeed co-evolution, of the supporting stroma. We assumed simple relationships between genotype, phenotype and fitness, and also neglected to model cell-cell interactions. Critically, we also used an abstract fitness function to define cellular phenotypes, and in doing so neglected to describe any molecular details of cell behaviour. Adequately describing these kinds of important biological complexities within a model is a necessary next step for the development of *in silico* biomarker development platform that is of general use. Increasing the realism of the model would improve confidence that the predicted prognostic value of any biomarker was not an artefact of the over-simplified model, although we have shown that our results are somewhat robust to alterations of a number of the key parameters in our model. Incorporating additional biological realism would also facilitate the *in silico* testing of the prognostic value of a full range of specific biological features; for example, the expression of a protein that fulfills a particular biological function, such as modulating cell adhesion.

Our study demonstrates how a computational model offers a platform for the initial development of novel prognostic biomarkers: computational models can be viewed as a high-throughput and cost-effective screening tool with which to identify the most promising biomarkers for subsequent empirical testing. This work provides the rationale for constructing an *in silico* biomarker development platform that would lessen the current restrictions imposed by the sole reliance on empirical testing.

## Acknowledgments

AD is supported by a J.D. Hatcher Award, School of Medicine, Queen’s University, Canada. AGF acknowledges support by the EPSRC through grant EP/I017909/1 (www.2020science.net). TAG is supported by Cancer Research UK. The authors wish to thank the anonymous reviewers for their insightful comments and suggested improvements to the manuscript.

## Text S1: Supplemental material

This document provides further details on the software implementation of the model described in the main text, as well as a detailed investigation of the robustness of the model to parameter values and constitutive assumptions.

**Software implementation.** A zipped folder is provided that contains the MATLAB implementation of the computational model and analysis described in this study. This folder includes a thoroughly documented example of how to run the code in the file README.txt. The code is also available online at github.com/AlexFletcher/cancer-risk-biomarkers.git.

**Robustness of results to mutation rate,** *μ*. To determine the effects of the mutation rate, *μ*, on the statistical association of each biomarker to time to cancer, we considered the two cases *μ* = 0.05 and *μ* = 0.01, with all other parameters set to their default values. Hazards ratios are computed for these cases in Tables S2 and S3, respectively. We found that for smaller mutation rates, the strength of the correlation of each biomarker decreased, due to the longer timescale of cancer development (and so increased stochasticity). We note in particular that the IPP measure remained correlated in biopsy samples, even for the lowest value of *μ* considered.

**Robustness of results to biopsy size,** *N_b_*. To determine the effects of the biopsy size, *N_b_*, on the correlation of each biomarker with the to time to cancer, we considered the two cases *N_b_* = 5 and *N_b_* = 40, with all other parameters set to their default values. Hazards ratios are computed for these cases in Tables S4 and S5, respectively. As expected, when a larger biopsy is taken, the degree of correlation for each biomarker with the time to cancer increases. For the small biopsy (*N_b_* = 5) we found that no biomarker values achieved statistical significance, while for the large biopsy (*N_b_* = 40), the statistical association of the Gini-Simpson index and IPP measures reached statistical significance. These results suggest that there is a critical biopsy size at which biomarker values may gain statistical correlation and significance, and the sampling effects of noise can be overcome.

**Robustness of results to threshold for cancer,** *N_m_*. To examine the effects of the number of advantageous mutations necessary for a cell to be considered cancerous, *N_m_*, on the correlation of each biomarker with the waiting time to cancer, we considered the three cases *N_b_* ∈ {3, 5, 15}, with all other parameters set to their default values. Hazards ratios are computed for these cases in Tables S6, S7 and S8, respectively. In particular, we found that the diversity and IPP measures consistently provided a strong and statistically-significant hazards ratio when computed from the whole-lesion, but became non-significant for biopsies.

**Robustness of results to fitness changes due to mutation,** *s_p_* **and** *s_d_*. To examine the effects of the degree to which non-neutral mutations alters cellular fitness on the association of each biomarker with the to time to cancer, we considered the three cases *s_p_* = *s_d_* ∈ {0,0.002,0.02}, with all other parameters set to their default values. Hazards ratios are computed for these cases in Tables S9, S10 and S11. In particular, we found that as selective advantage or disadvantage varied, the clonal diversity and IPP measures remained correlated with outcome for whole-lesion sampling.

**Variations on proliferation indices.** We next explored further variations on the proliferation indices defined in the main text. First, we varied the number of advantageous mutations required to be considered a ‘positive cell’ by examining the cases defined as IPP 1 and IPP 3, where the *f_i_* in equation (8) are re-defined such that for IPP *k* we have 

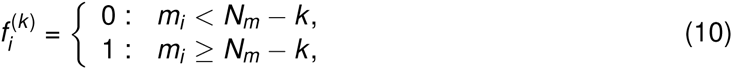
and the remainder of the formula for the IPP measure is unchanged. In addition, we consider a measure termed the index of mutation proliferation (IMP), defined analogously to the IPP, but with a choice of fitness function such that 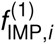 is the number of advantageous mutations obtained by cell *i* (IMP 1), and 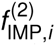 is the total number of mutations (the sum of advantageous, deleterious and neutral mutations) accumulated by cell *i* (IMP 2).

Results depicting the statistical association of these additional indices, as a function of sampling time, on both the whole-lesion and biopsy samples is summarised in Fig. S1. These results indicate that the behaviour of IPP1 and IPP3 is broadly similar to that of the IPP, albeit with their degrees of correlation peaking at different times.

**Effect of cell neighbourhood in a simplified model.** Having investigated the sensitivity of our results to the values of model parameters, we next examined whether the choice of cell neighbourhood has any significant effect on model behaviour. In lattice-based models such as our spatial Moran process, two common choices for defining the local topology are the Moore and von Neumann neighbourhoods. The von Neumann neighbours of a given cell are those that lie one lattice spacing from it by the Manhattan metric, while the Moore neighbours are those that lie less than two lattice spacings from it by the Euclidean metric.

To investigate whether the choice of neighbourhood has any impact on the behaviour of our mathematical model, we considered a simplified model with two cell types, which we refer to as normal and mutant. We employed the ‘death-birth’ update rule as defined in the main text. For simplicity we considered an irreversible neutral mutation giving no relative fitness advantage to mutants, with two cases of mutation probability considered: *μ* = 0 (corresponding to a classical spatial Moran process), and *μ* = 0.3. We considered a small 10 × 10 lattice comprised of cells and introduced a single mutant cell at the centre of the lattice. The model was simulated until the lattice was composed entirely of non-mutant cells or entirely of mutant cells (one of these possibilities must occur eventually, since they are both absorbing states of the model).

The cumulative distribution function for the time to fixation for the mutant cell population, computed based on 10^5^ runs of the above model using either Moore or von Neumann cell neighbourhoods, is shown in Fig. S2. We found that as the mutation rate is increased, the difference in the CDFs becomes nearly negligible. Additionally, in the case where *μ* = 0, for the von Neumann neighbourhood case, we found that 1.13% of the 10^5^ runs end in mutant fixation, and in the case of the Moore neighbourhood, 1.19% of the runs end in mutant fixation. When *μ* = 0.3, the probability of mutant fixation is higher, and for the von Neumann neighbourhood case, we find that 97.7% of the 2000 runs end in mutant fixation, whereas for the Moore neighbourhood the proportion is 98.2%. In both cases, the differences between the two cases of neighbourhood types diminished as *μ* increased, and limiting-time behaviour cases for both systems was highly similar.

**Robustness of results to update rules.** To examine the effect that the choice of update rule has on the behaviour of our model, we first considered 8 different update rules and compared key summary statistics in each case, for the simplified model described in the previous section. In this case, we incorporated an advantage to the mutations incurred by the mutant cells, conferring upon them a relative fitness of *s_p_* = 1.2. We use a two-letter notation to described the different choices of update rule, where the first and second letters represent the choice of first and second steps in the update rule, respectively. We use the letters *b* and *d* to denote birth (division) and death (removal), and we use a capital letter to denote the case where the corresponding step is influenced by cellular fitness, enabling the incorporation of selection. For instance, *bD* represents the update rule where first a cell is chosen uniformly at random to divide, and then a neighbouring cell is chosen to die, biased by inverse fitness.

We examined the average time to fixation for a mutant-only population, and the variance in these times to fixation, as well as the cumulative distribution function (CDF) for the time to fixation of the mutant population, conditional on this occurring before the mutants entirely die out. These results are presented for the case where *μ* = 0 in Table S12 and Fig. S3. We conclude that in this case, the various update rules produce differing results, but that as *μ* increases, these differences decrease in magnitude, as the times to fixation are reduced, and the probability of mutant fixation approaches unity. That is, the results presented in the above analysis depict upper bounds on the differences, and we note that based on our analysis, these differences decrease as *μ* is increased above zero, into the regimes of values considered within the present study, and those that are biologically relevant.

To further investigate how the prognostic value of the various candidate biomarkers depended on the choice of update rule, we performed additional model simulations where proliferation was now modified to be a birth-death (bD), rather than a death-birth (Db), process. In these simulations, a cell was chosen uniformly at random to divide, and its daughter cell (which, with some probability, accumulated a mutation) replaced a neighbour at random based on its fitness. The results of these simulations are shown in Table S13. In summary, we found that: (i) the time to cancer was markedly shortened compared to the death-birth case; (ii) the prognostic value of one of the putative biomarkers, the number of cells with at least two advantageous mutations, between significantly correlated with the waiting time to cancer; (iii) the mitotic proportion of cells remained a poor prognosticator; (iv) the clonal diversity measures (Simpson and Shannon indices) remained prognostic, and (v) the IPP and INP remained the strongest predictors of cancer risk among the putative biomarkers considered. We note also that the biopsy-based indices lost their association with the waiting time to cancer in the birth-death model.

**Robustness of results to decoupling mutation and proliferation.** A key assumption in our model was that cells mutate only when dividing. Thus, a cell that is adjacent to lower-fitness cells will accumulate mutations faster than a cell that is adjacent to higher-fitness cells, because the latter will have fewer opportunities to divide. To examine the robustness of our results to this assumption, we performed additional simulations in the case where mutations were decoupled from divisions. In these simulations, cell death and birth events were implemented as before, but all divisions were assumed to occur without mutation. Instead, immediately after each division, we drew a random number of mutations from a Poisson distribution with mean 0.5 (whose value is chosen arbitrarily to represent the mutation rate) and randomly bestowed each mutation on a cell in the lattice, independently chosen uniformly at random with replacement.

The results of these simulations are summarized in Table S14. We observed that decoupling mutation and division alters the predictive value of the various measures but that the general pattern of predictability is unaltered, with diversity measures remaining amongst the best predictors across the different tissue sampling methods. When spatial information is available (for whole lattice and biopsy measures), the IPP continues to perform well.

**Robustness of results to symmetry of mutations.** To determine how robust our results are to the key model assumption that mutations were equally likely to be advantageous, neutral or deleterious, we performed additional simulations where this symmetry was broken. In these simulations, the probabilities of a mutation being positive, neutral or deleterious was given by parameters *p*_adv_, *p*_neut_ and *p*_del_, which summed to 1 but need not be equal. Results of each of these simulations are summarized in Tables S15–S17.

Throughout each of these cases, we observed that the IPP and INP remained consistently statistically significant when the whole lesion is sampled, and the Shannon and Gini-Simpson indices are most correlated with the waiting time to cancer when the probability of an advantageous mutation is greatest. Furthermore, among each of these asymmetric cases for mutation probability, Moran’s *I* and Geary’s *C* were found to be extremely non-predictive of outcome. Overall, however, these results do not differ qualitatively from the symmetric case (*p*_adv_ = *p*_neut_ = *p*_del_ = 1/3) considered in the main text. Importantly, among each of these cases, the correlation between the IPP and INP measures and the waiting time remained strong and statistically significant. The results suggest a relative insensitivity of our biomarker results to this underlying model assumption.

**Tables**

**Table S1.** Summary of Cox proportional hazards models for varying values of the ‘positive staining’ threshold, *N_p_*. Hazard ratios (HR75) are computed at time *t* = 75 for the case *N_m_* = 10, *s_p_* = *s_d_* = 0.2, and *μ* = 0.1. Statistically significant values are in bold.

| | | Unit change | HR75 | 95% CI | $p$ |
| --- | --- | --- | --- | --- | --- |
| Whole lesion | $N_p = 2$ | 0.01 | 0 | $(0, \infty)$ | 0.29 |
| | $N_p = 3$ | 0.01 | 2.4 | $(0.5, 11)$ | 0.28 |
| | $N_p = 5$ | 0.01 | **1.2** | $(1.1, 1.3)$ | $< 10^{-4}$ |
| | $N_p = 7$ | 0.01 | **1.4** | $(1.3, 1.4)$ | $< 10^{-4}$ |
| | $N_p = 9$ | 0.01 | **3.5** | $(2.9, 4.3)$ | $< 10^{-4}$ |

**Table S2.**
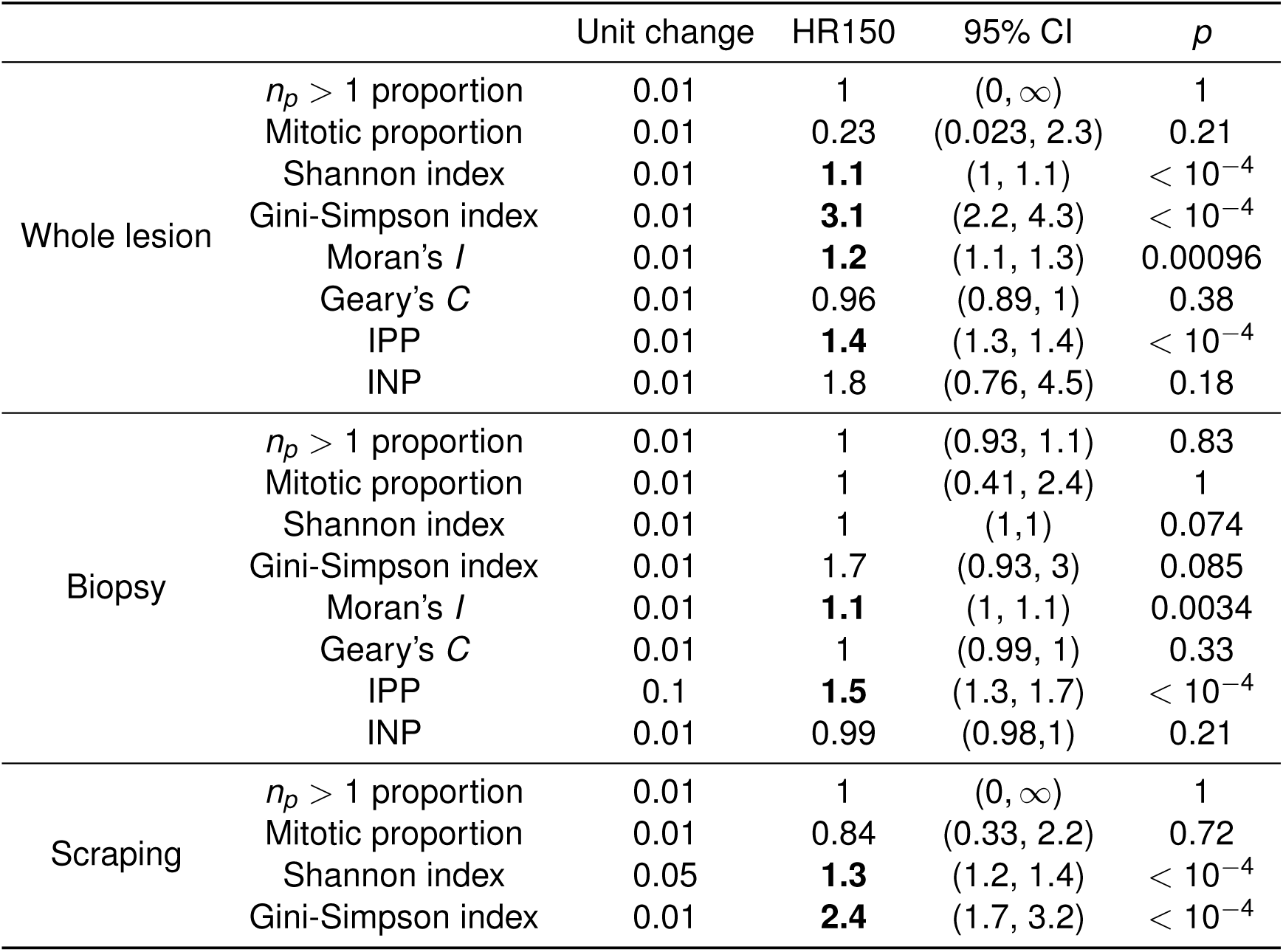
Summary of Cox proportional hazards models for the case of a lower mutation rate (*μ* = 0.005). Hazard ratios (HR150) are computed at time *t* = 150 for various putative biomarker schemes, for different tissue sampling schemes. Other parameter values used are *N_b_* = 20, *N_m_* = 10, *s_p_* = *s_d_* = 0.02 and those listed in Table 1. Statistically significant values are in bold.

**Table S3.**
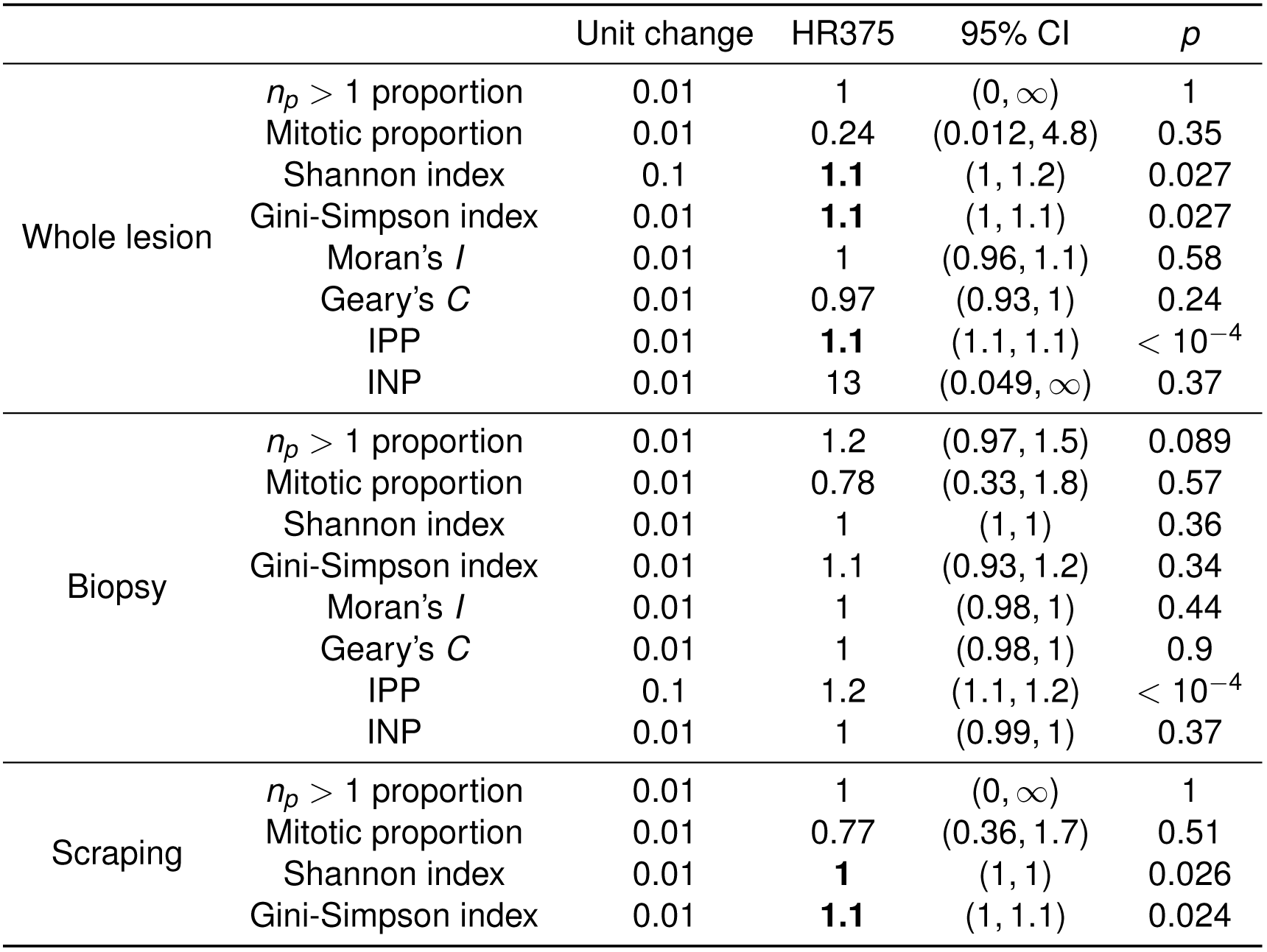
Summary of Cox proportional hazards models for the case of a much lower mutation rate (*μ* = 0.001). Hazard ratios (HR375) are computed at time *t* = 375 for various putative biomarker schemes, for different tissue sampling schemes. Other parameter values used are *N_b_* = 20, *N_m_* = 10, *s_p_* = *s_d_* = 0.02 and those listed in Table 1. Statistically significant values are in bold.

| | | Unit change | HR375 | 95% CI | $p$ |
| --- | --- | --- | --- | --- | --- |
| Whole lesion | $n_p > 1$ proportion | 0.01 | 1 | $(0, \infty)$ | 1 |
| | Mitotic proportion | 0.01 | 0.24 | $(0.012, 4.8)$ | 0.35 |
| | Shannon index | 0.1 | **1.1** | $(1, 1.2)$ | 0.027 |
| | Gini-Simpson index | 0.01 | **1.1** | $(1, 1.1)$ | 0.027 |
| | Moran's $I$ | 0.01 | 1 | $(0.96, 1.1)$ | 0.58 |
| | Geary's $C$ | 0.01 | 0.97 | $(0.93, 1)$ | 0.24 |
| | IPP | 0.01 | **1.1** | $(1.1, 1.1)$ | $< 10^{-4}$ |
| | INP | 0.01 | 13 | $(0.049, \infty)$ | 0.37 |
| Biopsy | $n_p > 1$ proportion | 0.01 | 1.2 | $(0.97, 1.5)$ | 0.089 |
| | Mitotic proportion | 0.01 | 0.78 | $(0.33, 1.8)$ | 0.57 |
| | Shannon index | 0.01 | 1 | $(1, 1)$ | 0.36 |
| | Gini-Simpson index | 0.01 | 1.1 | $(0.93, 1.2)$ | 0.34 |
| | Moran's $I$ | 0.01 | 1 | $(0.98, 1)$ | 0.44 |
| | Geary's $C$ | 0.01 | 1 | $(0.98, 1)$ | 0.9 |
| | IPP | 0.1 | 1.2 | $(1.1, 1.2)$ | $< 10^{-4}$ |
| | INP | 0.01 | 1 | $(0.99, 1)$ | 0.37 |
| Scraping | $n_p > 1$ proportion | 0.01 | 1 | $(0, \infty)$ | 1 |
| | Mitotic proportion | 0.01 | 0.77 | $(0.36, 1.7)$ | 0.51 |
| | Shannon index | 0.01 | **1** | $(1, 1)$ | 0.026 |
| | Gini-Simpson index | 0.01 | **1.1** | $(1, 1.1)$ | 0.024 |

**Table S4.** Summary of Cox proportional hazards models for the case of a smaller biopsy (*N_b_* = 5). Hazard ratios (HR75) are computed at time *t* = 75 for various putative biomarkers, for a circular biopsy. Other parameter values used are *N_m_* = 10, *s_p_* = *s_d_* = 0.2, *μ* = 0.1 and those listed in Table 1. Statistically significant values are in bold.

| | | Unit change | HR75 | 95% CI | $p$ |
| --- | --- | --- | --- | --- | --- |
| Biopsy | $n_p > 1$ proportion | 0.01 | 1 | (0.98, 1) | 0.75 |
|  | Mitotic proportion | 0.01 | 1 | (0.87, 1.2) | 0.67 |
|  | Shannon index | 0.01 | 1 | (0.99, 1) | 0.27 |
|  | Gini-Simpson index | 0.01 | 0.94 | (0.84, 1.1) | 0.3 |
| | Moran's $I$ | 0.01 | 1 | (0.96, 1) | 0.93 |
| | Geary's $C$ | 0.01 | 1 | (0.99, 1) | 0.44 |
|  | IPP | 0.01 | 1 | (0.97, 1) | 0.81 |
|  | INP | 0.01 | 1 | (1, 1) | 0.95 |

**Table S5.** Summary of Cox proportional hazards models for the case of a larger biopsy (*N_b_* = 40). Hazard ratios (HR12) are computed at time *t* = 12 for various putative biomarker schemes, for a circular biopsy. Other parameter values used are *N_m_* = 10, *s_p_* = *s_d_* = 0.2, *μ* = 0.1 and those listed in Table 1. Statistically significant values are in bold.

| | | Unit change | HR75 | 95% CI | $p$ |
| --- | --- | --- | --- | --- | --- |
| Biopsy | $n_p > 1$ proportion | 0.01 | 1 | (0.92, 1.2) | 0.6 |
|  | Mitotic proportion | 0.01 | 2.1 | (0.46, 9.6) | 0.34 |
|  | Shannon index | 0.1 | 1.2 | (1, 1.5) | 0.018 |
|  | Gini-Simpson index | 0.01 | **11** | (1.5, 87) | 0.018 |
| | Moran's $I$ | 0.01 | 1 | (0.9, 1.2) | 0.67 |
| | Geary's $C$ | 0.01 | 1 | (0.97, 1) | 0.82 |
| | IPP | 0.01 | **1.3** | (1.2, 1.4) | $< 10^{-4}$ |
|  | INP | 0.01 | 1.1 | (0.96, 1.2) | 0.19 |

**Table S6.** Summary of Cox proportional hazards models for the case of a much lower threshold number of mutations for cancer (*N_m_* = 3). Hazard ratios (HR12) are computed at time *t* = 12 for various putative biomarker schemes, for different tissue sampling schemes. Other parameter values used are *N_b_* = 20, *s_p_* = *s_d_* = 0.2, *μ* = 0.1 and those listed in Table 1. Statistically significant values are in bold.

| | | Unit change | HR12 | 95% CI | $p$ |
| --- | --- | --- | --- | --- | --- |
| Whole lesion | $n_p > 1$ proportion | 0.01 | **2.4** | (2.1, 2.8) | $< 10^{-4}$ |
|  | Mitotic proportion | 0.01 | 1.3 | (0.27, 6.3) | 0.73 |
| | Shannon index | 0.01 | **1.2** | (1.1, 1.2) | $< 10^{-4}$ |
| | Gini-Simpson index | 0.01 | **2.7** | (2, 3.7) | $< 10^{-4}$ |
| | Moran's $I$ | 0.01 | **6.6** | (2.1, 20) | 0.0011 |
| | Geary's $C$ | 0.01 | 1 | (0.8, 1.3) | 0.9 |
| | IPP | 0.01 | **1.3** | (1.2, 1.5) | $< 10^{-4}$ |
| | INP | 0.01 | **1.4** | (1.2, 1.5) | $< 10^{-4}$ |
| Biopsy | $n_p > 1$ proportion | 0.01 | 1.1 | (0.87, 1.5) | 0.37 |
|  | Mitotic proportion | 0.01 | 0.73 | (0.43, 1.2) | 0.25 |
|  | Shannon index | 0.01 | 1 | (0.99, 1) | 0.62 |
|  | Gini-Simpson index | 0.01 | 1.1 | (0.81, 1.6) | 0.49 |
| | Moran's $I$ | 0.01 | 1.1 | (0.88, 1.4) | 0.4 |
| | Geary's $C$ | 0.01 | 1 | (0.98, 1) | 0.74 |
|  | IPP | 0.1 | 1.6 | (1.2, 2.1) | 0.0028 |
|  | INP | 0.1 | 1.6 | (1.1, 2.3) | 0.0083 |
| Scraping | $n_p > 1$ proportion | 0.01 | **1.4** | (1.3, 1.5) | $< 10^{-4}$ |
|  | Mitotic proportion | 0.01 | 1.4 | (0.92, 2.2) | 0.11 |
|  | Shannon index | 0.1 | 1.5 | (1.1, 2.1) | 0.0049 |
|  | Gini-Simpson index | 0.01 | **1.3** | (1, 1.5) | 0.015 |

**Table S7.**
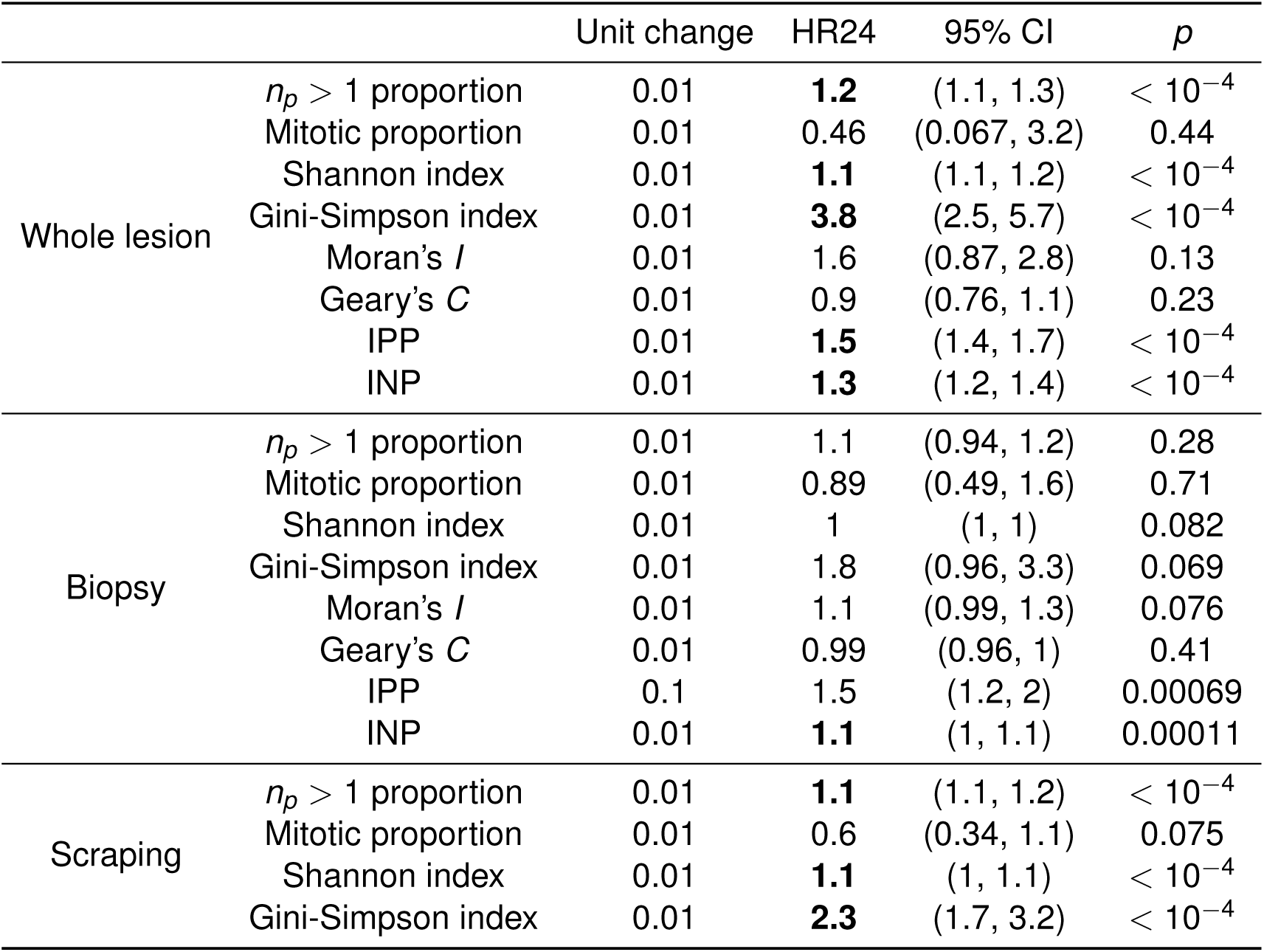
Summary of Cox proportional hazards models for the case of a lower threshold number of mutations for cancer (*N_m_* = 5). Hazard ratios (HR24) are computed at time *t* = 24 for various putative biomarker schemes, for different tissue sampling schemes. Other parameter values used are *N_b_* = 20, *s_p_* = *s_d_* = 0.2, *μ* = 0.1 and those listed in Table 1. Statistically significant values are in bold.

| | | Unit change | HR24 | 95% CI | $p$ |
| --- | --- | --- | --- | --- | --- |
| Whole lesion | $n_p > 1$ proportion | 0.01 | **1.2** | (1.1, 1.3) | $< 10^{-4}$ |
|  | Mitotic proportion | 0.01 | 0.46 | (0.067, 3.2) | 0.44 |
| | Shannon index | 0.01 | **1.1** | (1.1, 1.2) | $< 10^{-4}$ |
| | Gini-Simpson index | 0.01 | **3.8** | (2.5, 5.7) | $< 10^{-4}$ |
| | Moran's $I$ | 0.01 | 1.6 | (0.87, 2.8) | 0.13 |
| | Geary's $C$ | 0.01 | 0.9 | (0.76, 1.1) | 0.23 |
| | IPP | 0.01 | **1.5** | (1.4, 1.7) | $< 10^{-4}$ |
| | INP | 0.01 | **1.3** | (1.2, 1.4) | $< 10^{-4}$ |
| Biopsy | $n_p > 1$ proportion | 0.01 | 1.1 | (0.94, 1.2) | 0.28 |
|  | Mitotic proportion | 0.01 | 0.89 | (0.49, 1.6) | 0.71 |
|  | Shannon index | 0.01 | 1 | (1, 1) | 0.082 |
|  | Gini-Simpson index | 0.01 | 1.8 | (0.96, 3.3) | 0.069 |
| | Moran's $I$ | 0.01 | 1.1 | (0.99, 1.3) | 0.076 |
| | Geary's $C$ | 0.01 | 0.99 | (0.96, 1) | 0.41 |
|  | IPP | 0.1 | 1.5 | (1.2, 2) | 0.00069 |
|  | INP | 0.01 | **1.1** | (1, 1.1) | 0.00011 |
| Scraping | $n_p > 1$ proportion | 0.01 | **1.1** | (1.1, 1.2) | $< 10^{-4}$ |
|  | Mitotic proportion | 0.01 | 0.6 | (0.34, 1.1) | 0.075 |
| | Shannon index | 0.01 | **1.1** | (1, 1.1) | $< 10^{-4}$ |
| | Gini-Simpson index | 0.01 | **2.3** | (1.7, 3.2) | $< 10^{-4}$ |

**Table S8.** Summary of Cox proportional hazards models for the case of a larger threshold number of mutations for cancer (*N_m_* = 15). Hazard ratios (HR225) are computed at time *t* = 225 for various putative biomarker schemes, for different tissue sampling schemes. Other parameter values used are *N_b_* = 20, *s_p_* = *s_d_* = 0.2, *μ* = 0.1 and those listed in Table 1. Statistically significant values are in bold.

| | | Unit change | HR225 | 95% CI | $p$ |
| --- | --- | --- | --- | --- | --- |
| Whole lesion | $n_p > 1$ proportion | 0.01 | 1 | (0, ∞) | 1 |
|  | Mitotic proportion | 0.01 | **0.015** | (0, 0.51) | 0.019 |
| | Shannon index | 0.01 | **1.1** | (1.1, 1.1) | $< 10^{-4}$ |
| | Gini-Simpson index | 0.01 | **13** | (5.8, 29) | $< 10^{-4}$ |
| | Moran's $I$ | 0.01 | **1.3** | (1.1, 1.5) | 0.00079 |
| | Geary's $C$ | 0.01 | 0.91 | (0.83, 1) | 0.051 |
| | IPP | 0.01 | **1.7** | (1.6, 1.8) | $< 10^{-4}$ |
| | INP | 0.01 | 0.0059 | (0, ∞) | 0.93 |
| Biopsy | $n_p > 1$ proportion | 0.01 | 1 | (0.96, 1) | 0.77 |
|  | Mitotic proportion | 0.01 | 1.1 | (0.44, 2.8) | 0.83 |
|  | Shannon index | 0.01 | 1 | (1, 1) | 0.067 |
|  | Gini-Simpson index | 0.01 | 3.2 | (1, 11) | 0.051 |
| | Moran's $I$ | 0.01 | 0.99 | (0.95, 1) | 0.58 |
| | Geary's $C$ | 0.01 | 0.99 | (0.98, 1) | 0.57 |
|  | IPP | 0.1 | 1.4 | (1.2, 1.8) | 0.00013 |
|  | INP | 0.01 | 1 | (1, 1) | 0.67 |
| Scraping | $n_p > 1$ proportion | 0.01 | 1 | (0, ∞) | 1 |
|  | Mitotic proportion | 0.01 | 0.64 | (0.23, 1.8) | 0.4 |
| | Shannon index | 0.01 | **1.1** | (1, 1.1) | $< 10^{-4}$ |
| | Gini-Simpson index | 0.01 | **6.7** | (3.4, 13) | $< 10^{-4}$ |

**Table S9.** Summary of Cox proportional hazards models for the case where mutations do not alter cellular fitness (*s_p_* = *s_d_* = 0). Hazard ratios (HR105) are computed at time *t* = 105 for various putative biomarker schemes, for different tissue sampling schemes. Other parameter values used are *N_b_* = 20, *N_m_* = 10, *μ* = 0.1 and those listed in Table 1. Statistically significant values are in bold.

| | | Unit change | HR105 | 95% CI | $p$ |
| --- | --- | --- | --- | --- | --- |
| Whole lesion | $n_p > 1$ proportion | 0.01 | **1.1** | (1, 1.2) | 0.0022 |
|  | Mitotic proportion | 0.01 | 1.2 | (0.26, 5.2) | 0.85 |
| | Shannon index | 0.01 | **1.1** | (1, 1.1) | $< 10^{-4}$ |
|  | Gini-Simpson index | 0.001 | **1.9** | (1.2, 2.9) | 0.0052 |
| | Moran's $I$ | 0.01 | 0.83 | (0.64, 1.1) | 0.16 |
| | Geary's $C$ | 0.01 | 1.1 | (0.97, 1.2) | 0.15 |
| | IPP | 0.01 | **3.5** | (2.8, 4.4) | $< 10^{-4}$ |
|  | INP | 0.01 | **1.1** | (1, 1.1) | 0.014 |
| Biopsy | $n_p > 1$ proportion | 0.01 | 1 | (0.99, 1) | 0.55 |
|  | Mitotic proportion | 0.01 | 0.99 | (0.63, 1.6) | 0.98 |
|  | Shannon index | 0.01 | 1.4 | (1.2, 1.8) | 0.054 |
|  | Gini-Simpson index | 0.001 | 1.3 | (0.95, 1.8) | 0.1 |
| | Moran's $I$ | 0.01 | 1 | (0.92, 1.1) | 1 |
| | Geary's $C$ | 0.01 | 1 | (0.99, 1) | 0.34 |
| | IPP | 0.01 | **1.1** | (1.1, 1.2) | $< 10^{-4}$ |
|  | INP | 0.01 | 1 | (0.99, 1) | 0.16 |
| Scraping | $n_p > 1$ proportion | 0.01 | **1.1** | (1.1, 1.2) | 0.0003 |
|  | Mitotic proportion | 0.01 | 1 | (0.67, 1.5) | 0.98 |
|  | Shannon index | 0.1 | 1.5 | (1.2, 1.9) | 0.0019 |
|  | Gini-Simpson index | 0.001 | **1.6** | (1.1, 2.4) | 0.009 |

**Table S10.**
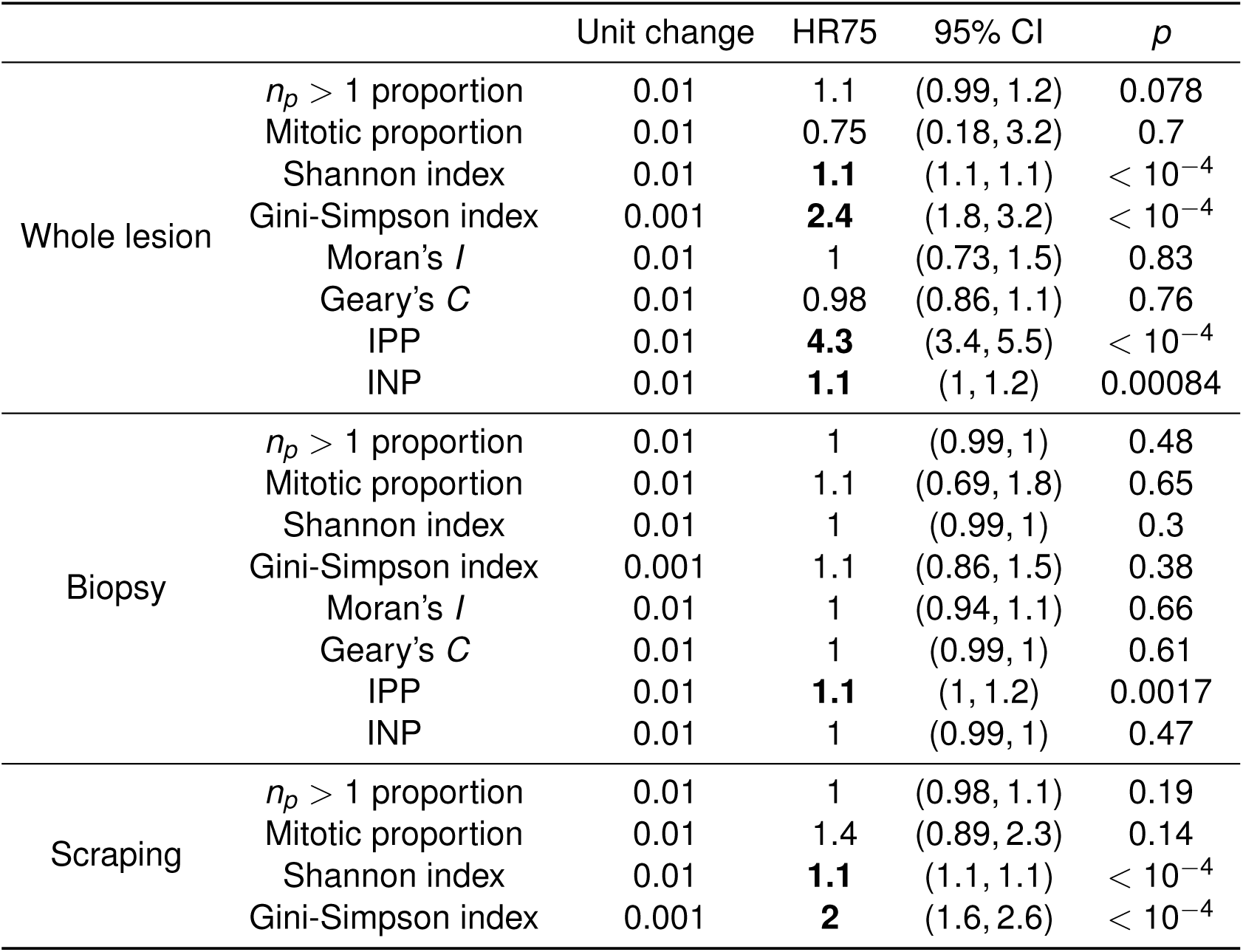
Summary of Cox proportional hazards models for the case where mutations alter cellular fitness to a lower extent (*s_p_* = *s_d_* = 0.002). Hazard ratios (HR75) are computed at time *t* = 75 for various putative biomarker schemes, for different tissue sampling schemes. Other parameter values used are *N_b_* = 20, *N_m_* = 10, *μ* = 0.1 and those listed in Table 1. Statistically significant values are in bold.

**Table S11.**
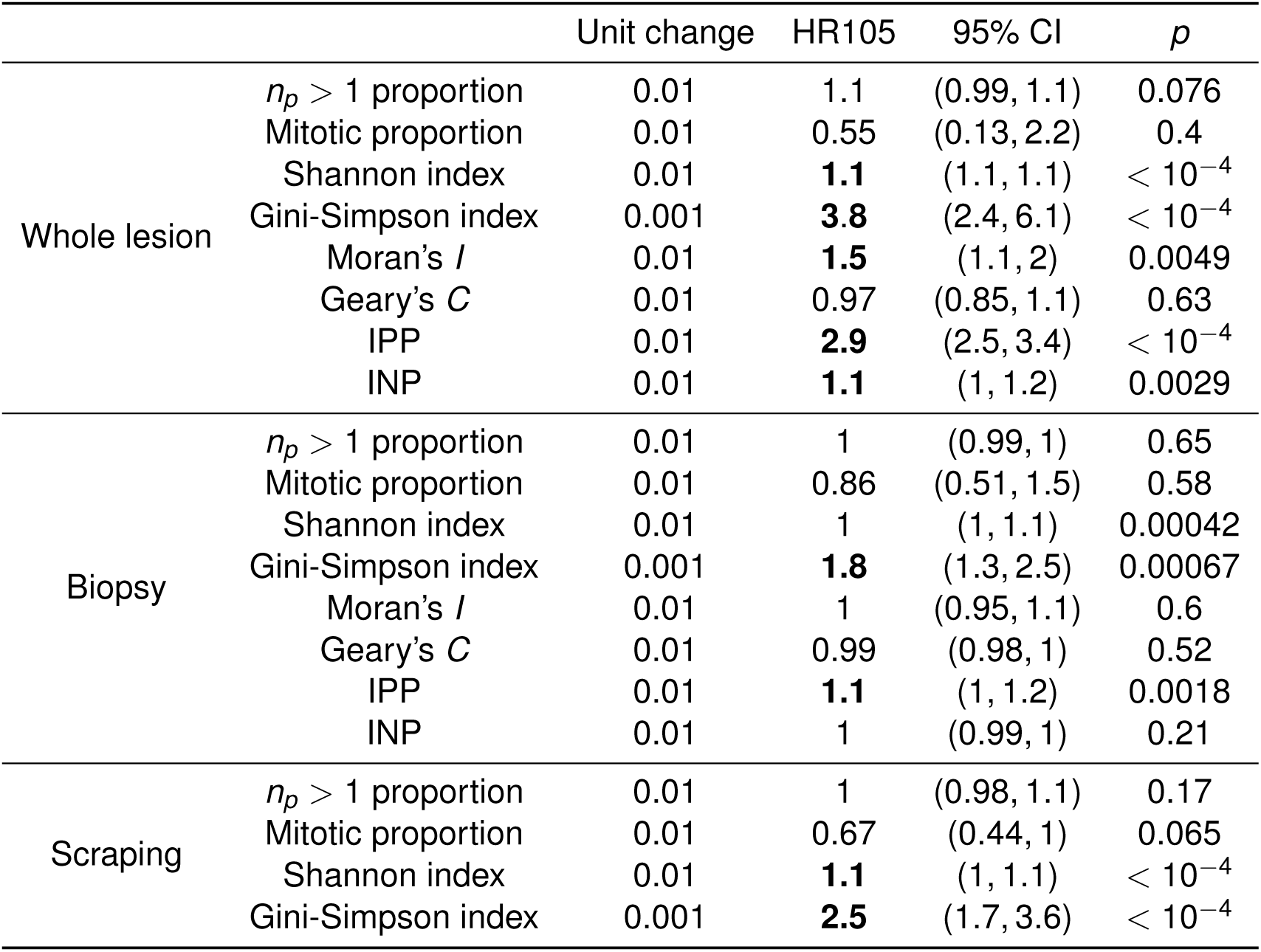
Summary of Cox proportional hazards models for the case where mutations alter cellular fitness to a much lower extent (*s_p_* = *s_d_* = 0.0002). Hazard ratios (HR105) are computed at time *t* = 105 for various putative biomarker schemes, for different tissue sampling schemes. Other parameter values used are *N_b_* = 20, *N_m_* = 10, *μ* = 0.1 and those listed in Table 1. Statistically significant values are in bold.

**Table S12.** Effect of update rule on mutant fixation statistics in a spatial Moran model with selection only. Summarized statistics from 10^5^ runs of each update rule used in the simplified Moran model as described in the Supplementary Text, in the case *μ* = 0. Proportion: the proportion of simulations in which mutant fixation occurred. Mean, variance: the mean and variance of the times to mutant fixation, across simulations in which this occurred. Simulations were run until the population consisted entirely of either non-mutant cells or mutant cells.

| Update rule | Proportion | Mean | Variance |
| --- | --- | --- | --- |
| db | 0.011 | 206 | 14144 |
| dB | 0.162 | 72.3 | 614 |
| Db | 0.170 | 78.7 | 700 |
| DB | 0.296 | 44.8 | 146 |
| bd | 0.009 | 244 | 19390 |
| bD | 0.154 | 81.3 | 719 |
| Bd | 0.162 | 73 | 595 |
| BD | 0.294 | 41.4 | 121 |

**Table S13.** Summary of Cox proportional hazards models for the case where division precedes removal. Hazard ratios (HR7) are computed at time *t* = 7. Parameter values used are *N_m_* = 10, *s_p_* = *s_d_* = 0.2, and *μ* = 0.1. Statistically significant values are in bold.

| | | Unit change | HR7 | 95% CI | $p$ |
| --- | --- | --- | --- | --- | --- |
| Whole lesion | $n_p > 1$ proportion | 0.01 | **1.3** | (1.1, 1.6) | 0.002 |
|  | Mitotic proportion | 0.01 | 0.68 | (0.17, 2.7) | 0.59 |
|  | Shannon index | 0.01 | **1.1** | (1, 1.1) | 0.00046 |
|  | Gini-Simpson index | 0.01 | **1.2** | (1, 1.3) | 0.04 |
| | Moran's $I$ | 0.01 | **1** | (1, 1) | 0.0097 |
| | Geary's $C$ | 0.01 | 1 | (1, 1) | 0.36 |
|  | IPP | 0.0001 | **8.9** | (2.8, 28) | 0.00024 |
| | INP | 0.0001 | **1.4** | (1.2, 1.6) | $< 10^{-4}$ |
| Biopsy | $n_p > 1$ proportion | 0.01 | 1.4 | (0.95, 2.2) | 0.088 |
|  | Mitotic proportion | 0.01 | 1.2 | (0.7, 1.9) | 0.57 |
|  | Shannon index | 0.01 | 1 | (0.99, 1) | 0.44 |
|  | Gini-Simpson index | 0.01 | 1.1 | (0.89, 1.4) | 0.34 |
| | Moran's $I$ | 0.01 | 1.2 | (0.89, 1.6) | 0.23 |
| | Geary's $C$ | 0.01 | 0.99 | (0.97, 1) | 0.59 |
| | IPP | 0.0001 | 1 | (0, $10^6$) | 1 |
| | INP | 0.0001 | 1 | (0, $10^6$) | 1 |
| Scraping | Shannon index | 0.01 | **1** | (1, 1.1) | 0.0045 |
|  | Gini-Simpson index | 0.01 | 1.1 | (0.98, 1.2) | 0.13 |
| | $n_p > 1$ proportion | 0.01 | **1.2** | (1.1, 1.4) | 0.002 |
|  | Mitotic proportion | 0.01 | 0.86 | (0.56, 1.3) | 0.5 |

**Table S14.**
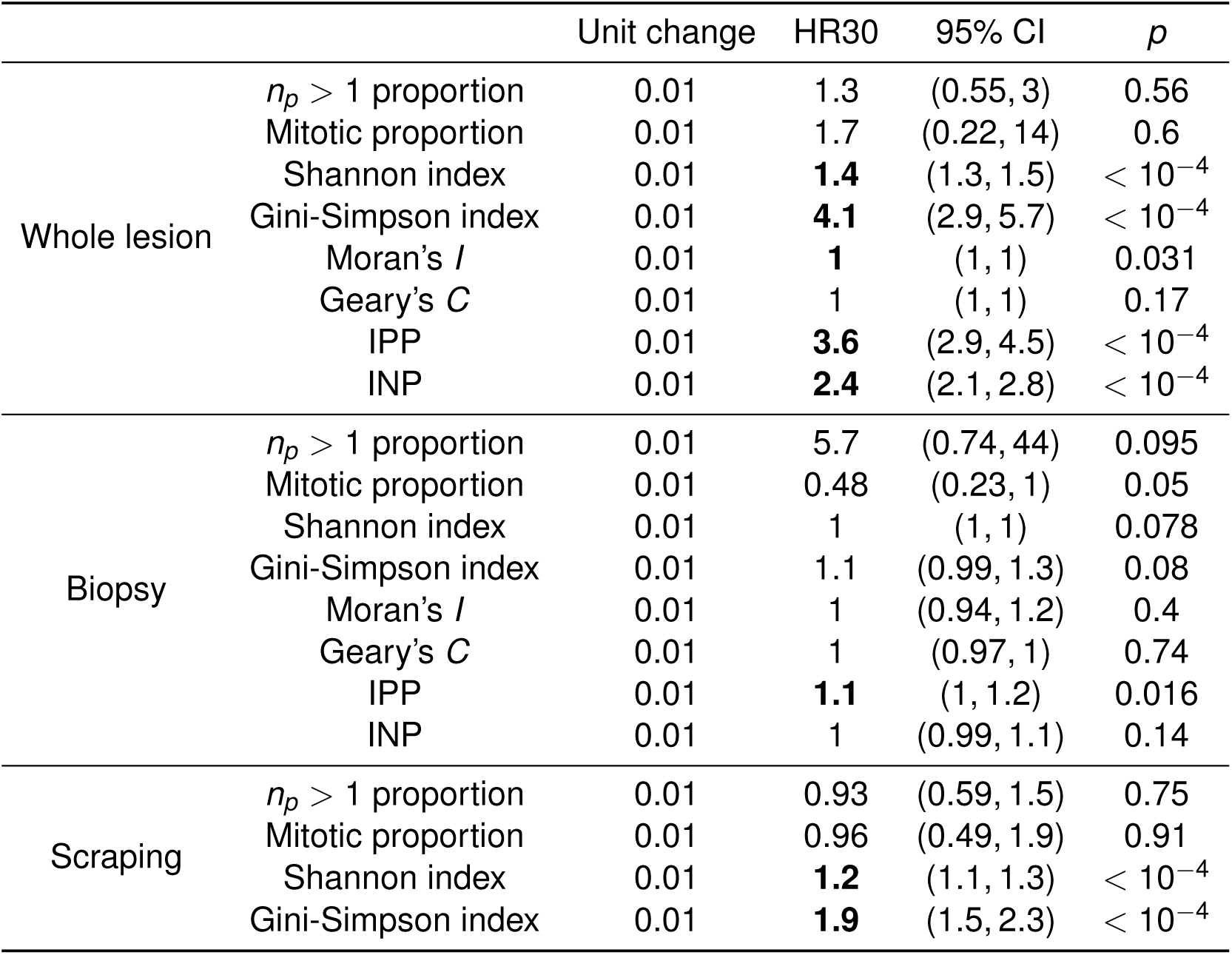
Summary of Cox proportional hazards models for the case where mutation and proliferation are decoupled. Hazard ratios (HR30) are computed at time *t* = 30. Parameter values used are *N_m_* = 10, *s_p_* = *s_d_* = 0.2 and *μ* = 0.1. A number of mutations drawn from the Poisson distribution with mean 0.5 are assumed to occur at each cell division, with each mutation bestowed independently upon a cell drawn uniformly at random from the lattice, with replacement. Statistically significant values are in bold.

| | | Unit change | HR30 | 95% CI | $p$ |
| --- | --- | --- | --- | --- | --- |
| Whole lesion | $n_p > 1$ proportion | 0.01 | 1.3 | (0.55, 3) | 0.56 |
|  | Mitotic proportion | 0.01 | 1.7 | (0.22, 14) | 0.6 |
| | Shannon index | 0.01 | **1.4** | (1.3, 1.5) | $< 10^{-4}$ |
| | Gini-Simpson index | 0.01 | **4.1** | (2.9, 5.7) | $< 10^{-4}$ |
| | Moran's $I$ | 0.01 | **1** | (1, 1) | 0.031 |
| | Geary's $C$ | 0.01 | 1 | (1, 1) | 0.17 |
| | IPP | 0.01 | **3.6** | (2.9, 4.5) | $< 10^{-4}$ |
| | INP | 0.01 | **2.4** | (2.1, 2.8) | $< 10^{-4}$ |
| Biopsy | $n_p > 1$ proportion | 0.01 | 5.7 | (0.74, 44) | 0.095 |
|  | Mitotic proportion | 0.01 | 0.48 | (0.23, 1) | 0.05 |
|  | Shannon index | 0.01 | 1 | (1, 1) | 0.078 |
|  | Gini-Simpson index | 0.01 | 1.1 | (0.99, 1.3) | 0.08 |
| | Moran's $I$ | 0.01 | 1 | (0.94, 1.2) | 0.4 |
| | Geary's $C$ | 0.01 | 1 | (0.97, 1) | 0.74 |
|  | IPP | 0.01 | **1.1** | (1, 1.2) | 0.016 |
|  | INP | 0.01 | 1 | (0.99, 1.1) | 0.14 |
| Scraping | $n_p > 1$ proportion | 0.01 | 0.93 | (0.59, 1.5) | 0.75 |
|  | Mitotic proportion | 0.01 | 0.96 | (0.49, 1.9) | 0.91 |
| | Shannon index | 0.01 | **1.2** | (1.1, 1.3) | $< 10^{-4}$ |
| | Gini-Simpson index | 0.01 | **1.9** | (1.5, 2.3) | $< 10^{-4}$ |

**Table S15.** Summary of Cox proportional hazards models for the case where mutations are more likely to be advantageous. Hazard ratios (HR30) are computed at time *t* = 30 for the case *N_m_* = 10, *s_p_* = *s_d_* = 0.2, and *μ* = 0.1. Here the probabilities of a mutation being advantageous, neutral or deleterious are given by *p*_pos_ = 0.8, *p*_neut_ = 0.1 and *p*_del_ = 0.1, respectively. Statistically significant values are in bold.

| | | Unit change | HR7 | 95% CI | $p$ |
| --- | --- | --- | --- | --- | --- |
| Whole lesion | $n_p > 1$ proportion | 0.01 | 0.59 | (0.25, 1.4) | 0.21 |
|  | Mitotic proportion | 0.01 | 2.7 | (0.34, 21) | 0.35 |
| | Shannon index | 0.01 | **1.3** | (1.2, 1.4) | $< 10^{-4}$ |
| | Gini-Simpson index | 0.01 | **3.1** | (2.3, 4.2) | $< 10^{-4}$ |
| | Moran's $I$ | 0.01 | 1 | (1, 1) | 0.71 |
| | Geary's $C$ | 0.01 | 1 | (1, 1) | 0.21 |
| | IPP | 0.01 | **2.6** | (2.1, 3.3) | $< 10^{-4}$ |
| | INP | 0.01 | **2** | (1.7, 2.3) | $< 10^{-4}$ |
| Biopsy | $n_p > 1$ proportion | 0.01 | 2.8 | (0.33, 23) | 0.35 |
|  | Mitotic proportion | 0.01 | 1.1 | (0.51, 2.1) | 0.89 |
|  | Shannon index | 0.01 | 1 | (1, 1) | 0.19 |
|  | Gini-Simpson index | 0.01 | 1.1 | (0.95, 1.2) | 0.21 |
| | Moran's $I$ | 0.01 | 1.1 | (0.95, 1.2) | 0.27 |
| | Geary's $C$ | 0.01 | 0.99 | (0.96, 1) | 0.37 |
|  | IPP | 0.01 | 1.1 | (0.99, 1.2) | 0.07 |
|  | INP | 0.01 | **1.1** | (1, 1.1) | 0.03 |
| Scraping | $n_p > 1$ proportion | 0.01 | 1 | (0.59, 1.7) | 0.97 |
|  | Mitotic proportion | 0.01 | 1.9 | (0.89, 4.2) | 0.099 |
| | Shannon index | 0.01 | **1.1** | (1.1, 1.2) | $< 10^{-4}$ |
|  | Gini-Simpson index | 0.01 | **1.4** | (1.1, 1.8) | 0.0014 |

**Table S16.** Summary of Cox proportional hazards models for the case where mutations are more likely to be neutral. Hazard ratios (HR30) are computed at time *t* = 30 for the case *N_m_* = 10, *s_p_* = *s_d_* = 0.2, and *μ* = 0.1. Here the probabilities of a mutation being advantageous, neutral or deleterious are given by *p*_pos_ = 0.1, *p*_neut_ = 0.8 and *p*_del_ = 0.1, respectively. Statistically significant values are in bold.

| | | Unit change | HR7 | 95% CI | $p$ |
| --- | --- | --- | --- | --- | --- |
| Whole lesion | $n_p > 1$ proportion | 0.01 | 1.1 | (0.52, 2.4) | 0.8 |
|  | Mitotic proportion | 0.01 | 0.14 | (0.018, 1) | 0.055 |
| | Shannon index | 0.01 | **1.3** | (1.2, 1.3) | $< 10^{-4}$ |
| | Gini-Simpson index | 0.01 | **2.6** | (2, 3.5) | $< 10^{-4}$ |
| | Moran's $I$ | 0.01 | 1 | (1, 1) | 0.11 |
| | Geary's $C$ | 0.01 | 1 | (1, 1) | 0.99 |
| | IPP | 0.01 | **3.4** | (2.7, 4.4) | $< 10^{-4}$ |
| | INP | 0.01 | **2** | (1.7, 2.3) | $< 10^{-4}$ |
| Biopsy | $n_p > 1$ proportion | 0.01 | **10** | (1.2, 91) | 0.034 |
|  | Mitotic proportion | 0.01 | 0.85 | (0.43, 1.7) | 0.65 |
|  | Shannon index | 0.01 | **1** | (1, 1) | 0.021 |
|  | Gini-Simpson index | 0.01 | **1.2** | (1, 1.3) | 0.02 |
| | Moran's $I$ | 0.01 | 0.98 | (0.87, 1.1) | 0.74 |
| | Geary's $C$ | 0.01 | 1 | (0.97, 1) | 0.85 |
|  | IPP | 0.01 | **1.1** | (1.1, 1.2) | 0.00092 |
|  | INP | 0.01 | **1.1** | (1, 1.1) | 0.0063 |
| Scraping | Shannon index | 0.01 | **1.1** | (1.1, 1.2) | $< 10^{-4}$ |
| | Gini-Simpson index | 0.01 | **1.7** | (1.4, 2.2) | $< 10^{-4}$ |
| | $n_p > 1$ proportion | 0.01 | 0.95 | (0.6, 1.5) | 0.83 |
|  | Mitotic proportion | 0.01 | 0.73 | (0.41, 1.3) | 0.29 |

**Table S17.** Summary of Cox proportional hazards models for the case where mutations are more likely to be deleterious. Hazard ratios (HR30) are computed at time *t* = 30 for the case *N_m_* = 10, *s_p_* = *s_d_* = 0.2, and *μ* = 0.1. Here the probabilities of a mutation being advantageous, neutral or deleterious are given by *p*_pos_ = 0.1, *p*_neut_ = 0.1 and *p*_del_ = 0.8, respectively. Statistically significant values are in bold.

| | | Unit change | HR7 | 95% CI | $p$ |
| --- | --- | --- | --- | --- | --- |
| Whole lesion | $n_p > 1$ proportion | 0.01 | **3** | (1.3, 6.8) | 0.0079 |
|  | Mitotic proportion | 0.01 | 5.2 | (0.75, 36) | 0.096 |
| | Shannon index | 0.01 | **1.2** | (1.1, 1.3) | $< 10^{-4}$ |
| | Gini-Simpson index | 0.01 | **2.4** | (1.8, 3.2) | $< 10^{-4}$ |
| | Moran's $I$ | 0.01 | **1** | (1, 1) | 0.0031 |
| | Geary's $C$ | 0.01 | 1 | (1, 1) | 0.24 |
| | IPP | 0.01 | **3.2** | (2.5, 4.1) | $< 10^{-4}$ |
| | INP | 0.01 | **1.9** | (1.7, 2.2) | $< 10^{-4}$ |
| Biopsy | $n_p > 1$ proportion | 0.01 | 6.2 | (0.63, 60) | 0.12 |
|  | Mitotic proportion | 0.01 | 1.5 | (0.74, 2.9) | 0.27 |
|  | Shannon index | 0.01 | 1 | (1, 1) | 0.11 |
|  | Gini-Simpson index | 0.01 | 1.1 | (0.98, 1.3) | 0.1 |
| | Moran's $I$ | 0.01 | 0.95 | (0.84, 1.1) | 0.36 |
| | Geary's $C$ | 0.01 | 0.99 | (0.96, 1) | 0.29 |
|  | IPP | 0.01 | **1.1** | (1, 1.2) | 0.0033 |
|  | INP | 0.01 | **1.1** | (1, 1.1) | 0.00062 |
| Scraping | Shannon index | 0.01 | **1.1** | (1.1, 1.2) | $< 10^{-4}$ |
| | Gini-Simpson index | 0.01 | **1.6** | (1.3, 2) | $< 10^{-4}$ |
| | $n_p > 1$ proportion | 0.01 | 1.1 | (0.73, 1.8) | 0.57 |
|  | Mitotic proportion | 0.01 | 1.4 | (0.7, 2.7) | 0.35 |

**Figure S1.**
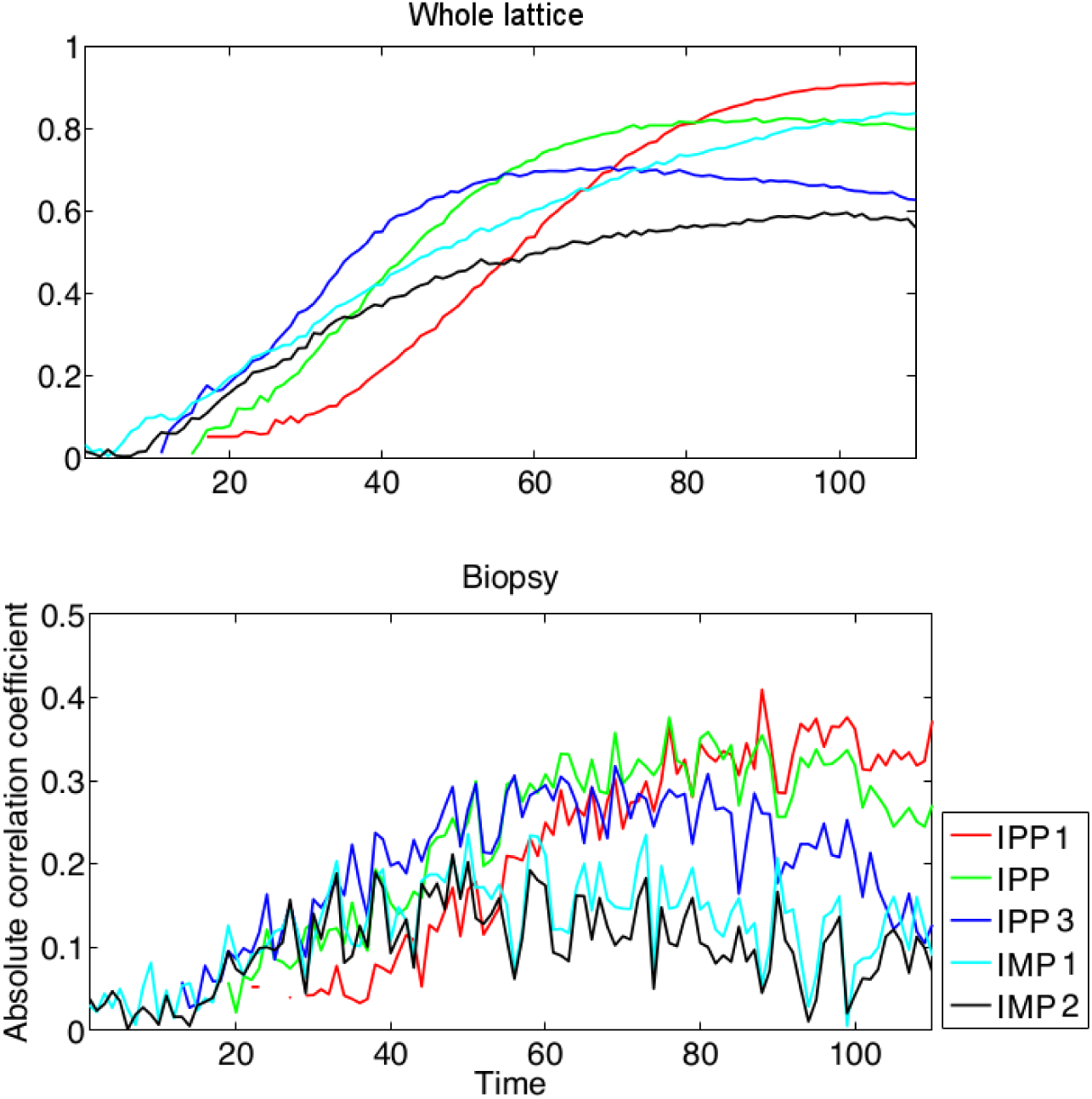
Variation of the absolute value of the correlation coefficient, as a function of sampling time *T_b_*, for additional indices considered in Text S1. Data shown are for the case where *N_m_* = 10, *s_p_* = *s_d_* = 0.2, and *μ* = 0.1, over 1000 runs, for indices taken on circular biopsies with radius *N_b_* = 20.

**Figure S2.**
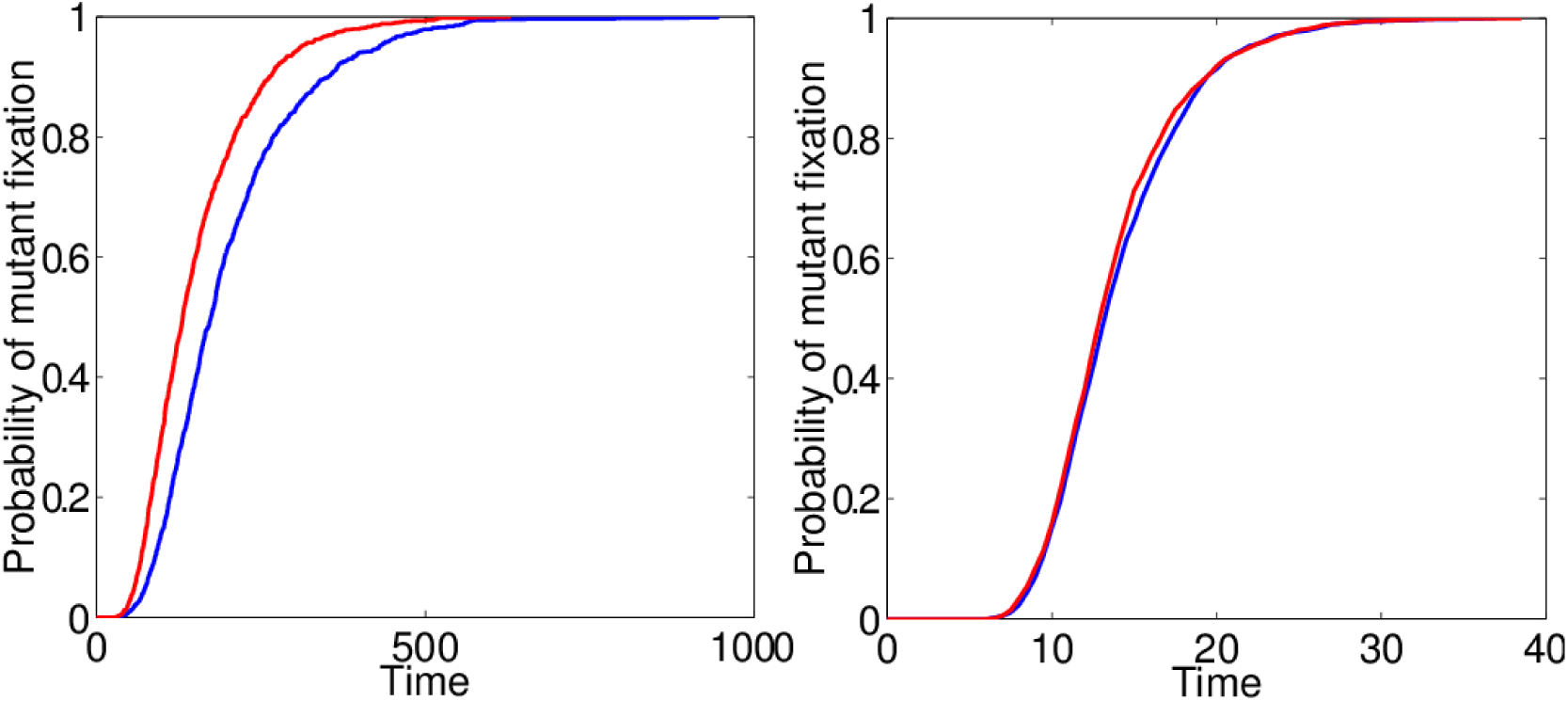
Cumulative distribution functions for the time to mutant fixation in a spatial Moran model with von Neumann and Moore neighbourhoods. The model includes a single, neutral, irreversible mutation (*s_p_* = *s_d_* = 0, *μ* = 0) and each simulation is initiated with a single mutant cell in the centre of the lattice. The blue and red curves correspond to the von Neumann and Moore neighbourhoods, respectively. Left: The case *μ* = 0; results from 10^5^ simulations. Right: The case *μ* = 0.3; results from 2000 simulations.

**Figure S3.**
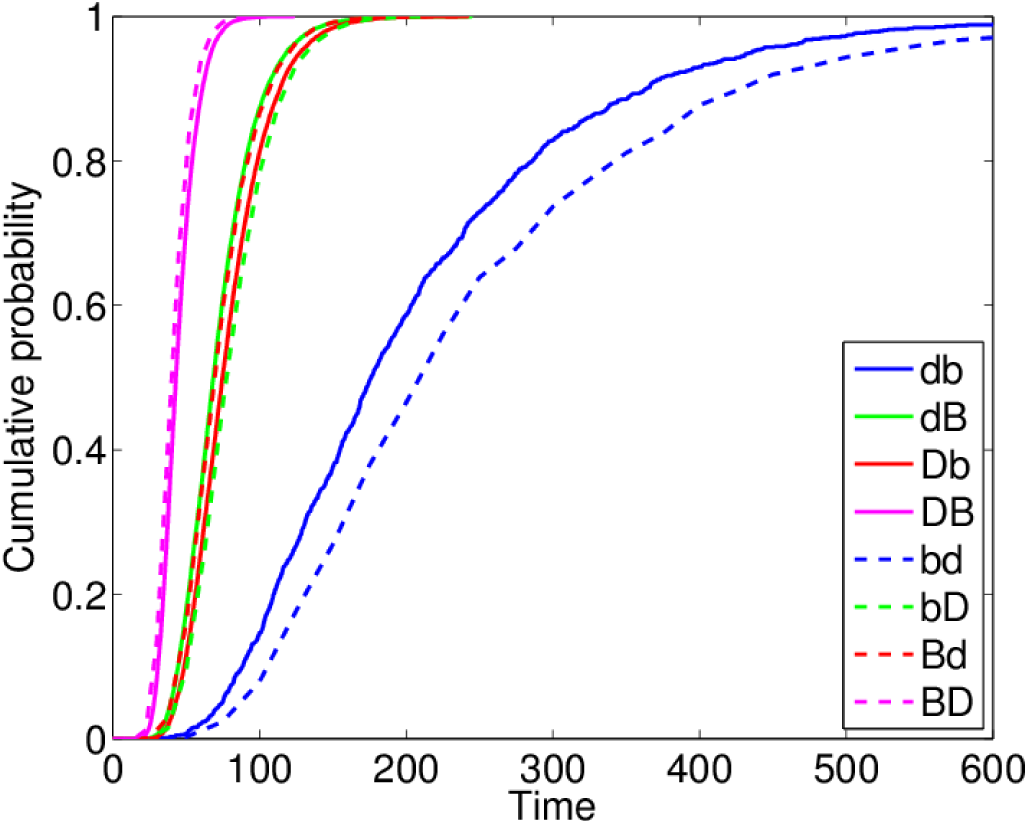
Cumulative distribution functions for the time to mutant fixation in a spatial Moran model under different update rules. The model includes a single, neutral, irreversible mutation (*sp* = *s_d_* = 0, *μ* = 0) and each simulation is initiated with a single mutant cell in the centre of the lattice. Legend: B and D correspond to birth and death respectively, written in the order in which these processes are implemented in the update rule, with capitalized letters indicating random selection of cells biased by their inverse fitness to account for the effects of selective advantage. Results are generated from 2000 simulations of a small 10 × 10 lattice.

**Figure S4.**
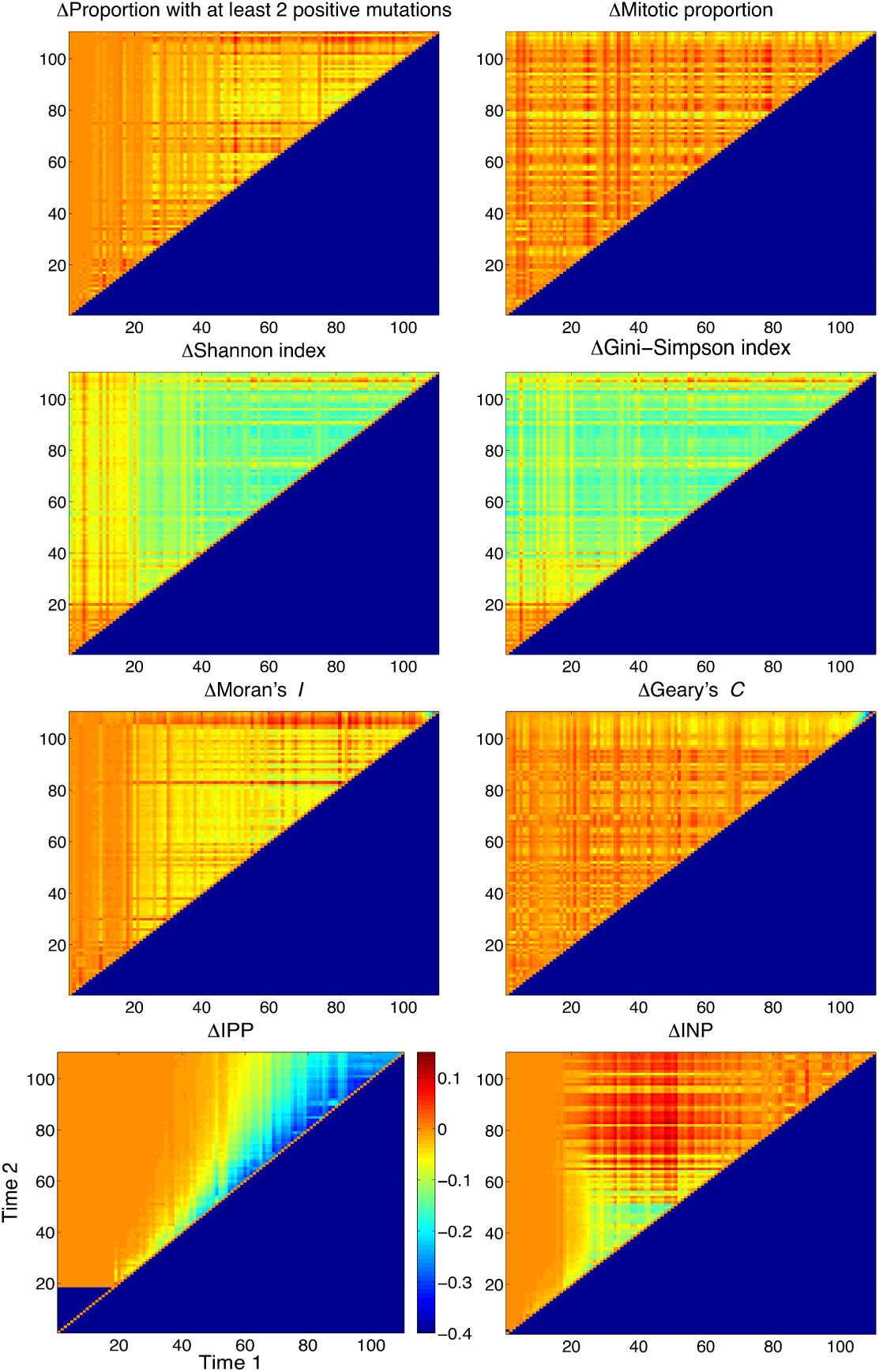
Taking the difference across serial biopsies provides only marginal additional prognostic information. Heat maps depicting the relative value of taking serial biopsies at different time points. Positive values (warm colours) indicate that prognostic value was improved by taking the difference between the biomarker value across the two time-points; negative values (cool colours) indicate that more information was available at the second time point alone. Results from 1000 simulations for each pair of time points, with *N_m_* = 10, *s_p_* = *s_d_* = 0.2, *μ* = 0.1 and *N_b_* = 20.

**Figure S5.**
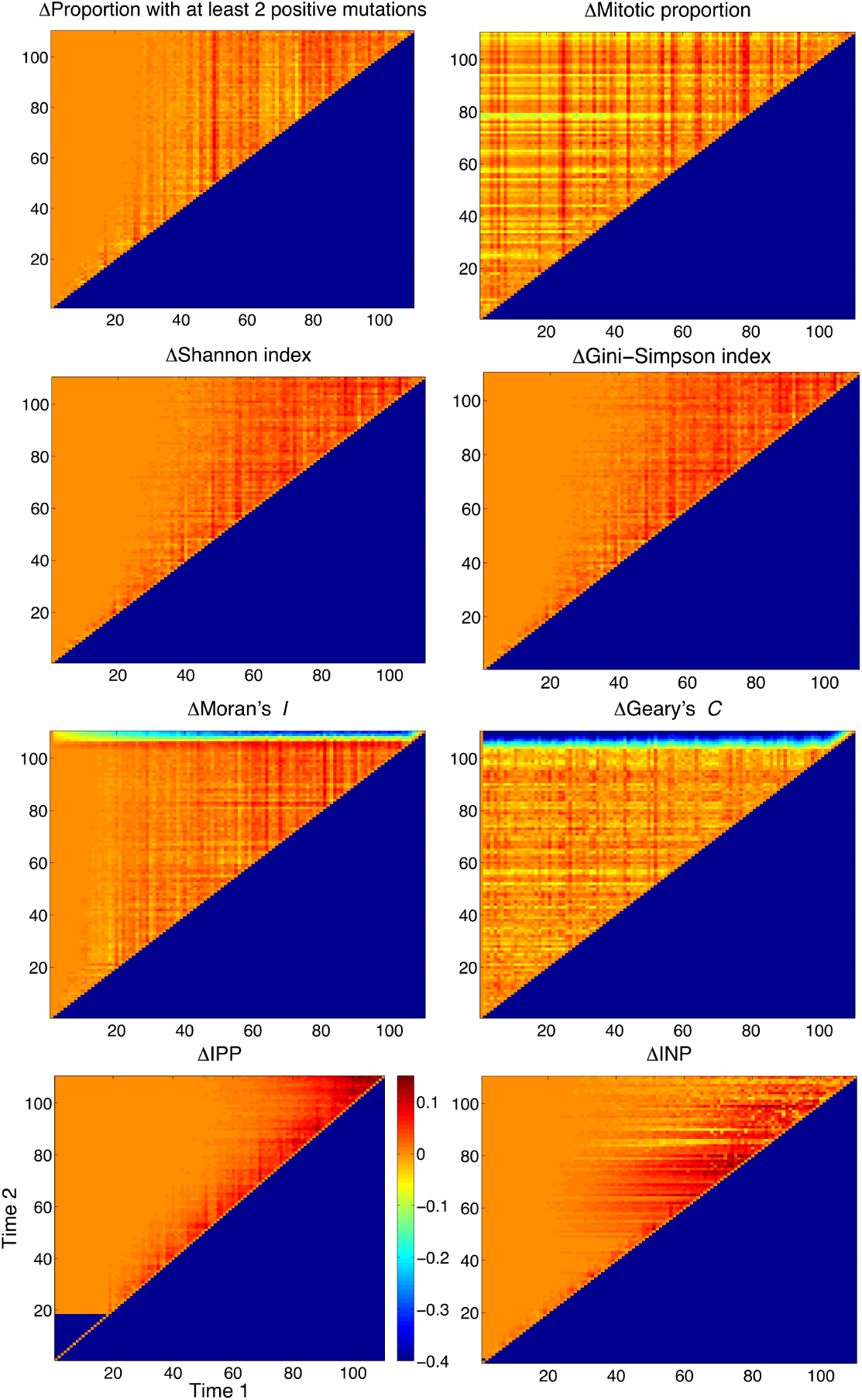
Maximizing biomarker values across serial biopsies provides only marginal additional prognostic information. Heat maps depicting the relative value of taking serial biopsies at different time points. Positive values (warm colours) indicate that prognostic value was improved by taking the maximum of the biomarker value across the two time-points; negative values (cool colours) indicate that more information was available at the second time point alone. Results from 1000 simulations for each pair of time points, with *N_m_* = 10, *s_p_* = *s_d_* = *0.2, μ* = 0.1 and *N_b_* = 20.

**Conflict of interest statement:** The authors declare no conflict of interest.

